# *Trachymyrmex septentrionalis* ant microbiome assembly is unique to individual colonies and castes

**DOI:** 10.1101/2021.11.04.467277

**Authors:** Emily A. Green, Jonathan L. Klassen

## Abstract

Within social insect colonies, microbiomes often differ between castes due to their different functional roles, and between colony locations. *Trachymyrmex septentrionalis* fungus-growing ants form colonies throughout the eastern USA and Northern Mexico that include workers, female and male alates (unmated reproductive castes), larvae, and pupae. How *T. septentrionalis* microbiomes vary across this geographic range and between castes is unknown. Our sampling of individual ants from colonies across the Eastern USA revealed a conserved *T. septentrionalis* worker ant microbiome, and that worker ant microbiomes are more conserved within colonies than between them. A deeper sampling of individual ants from two colonies that included all available castes (pupae, larvae, workers, female and male alates), from both before and after adaptation to controlled laboratory conditions, revealed that ant microbiomes from each colony, caste, and rearing condition were typically conserved within but not between each sampling category. Tenericute bacterial symbionts were especially abundant in these ant microbiomes and varied widely in abundance between sampling categories. This study demonstrates how individual insect colonies primarily drive the composition of their microbiomes, and that these microbiomes are further modified by developmental differences between insect castes and the different environmental conditions experienced by each colony.

**IMPORTANCE:** This study investigates microbiome assembly in the fungus-growing ant *Trachymyrmex septentrionalis*, showing how colony, caste, and lab adaptation influences the microbiome and revealing unique patterns of Mollicute symbiont abundance. We find that ant microbiomes differ strongly between colonies but less so within colonies. Microbiomes of different castes and following lab adaptation also differ in a colony-specific manner. This study advances understanding of the nature of individuality in social insect microbiomes, and cautions against the common practice of only sampling a limited number of populations to understand microbiome diversity and function.

## INTRODUCTION

Social insects live in colonies and manifest group integration, division of labor, and overlap of generations (1). These divisions of labor are often distributed between developmental castes, which at the most basic level are divided into reproductive individuals and sterile workers, which themselves can be further partitioned into specialized groups for different tasks. Gut microbiomes often differ between social insect castes, possibly due to the specialized functional role played by each caste (2). For example, the microbiomes of termite minor workers, major workers, and soldiers all differ from each other (3, 4), and termite reproductive castes have unique microbiomes compared to those of workers from the same colony (5–8). Honey bee queens, workers, and drones also each have unique gut microbiomes, where worker microbiomes are more diverse than those of queens and drones, possibly due to worker foraging (9). Honey bees have a well-defined core microbiome that is found in all colonies and castes, including queens, workers, and drones (10, 11). However, strains varied between geographic locations, individual colonies, and bee castes (9, 11). Social insect microbiome composition is therefore determined by a complex combination of developmental differences and colony-specific ecological factors.

Fungus-growing ants are social insects (Formicidae: tribe Attini) that evolved 55 to 65 MYA in South America (12–14). Today there are 19 genera including approximately 250 known species that range throughout North and South America from the northern USA to southern Argentina (14–18). The ∼250 species of fungus-growing ants are separated into five separate agricultural systems representing transitions in their evolution: lower, coral-fungus, yeast, higher, and leaf-cutter agriculture (14). Fungus-growing ant caste systems include workers, reproductive queens, and unmated reproductive alates. Leaf-cutting ants (*Atta, Acromyrmex*, and *Amiomyrmex*) can have multiple worker castes, including soldier, major, media and, minor workers (16). All fungus-growing ants grow a cultivar fungus, typically a species of *Leucoagaricus* (except for in coral-fungus agriculture by *Apterostigma* ants, which cultivates a Pterulaceae species (19)). This cultivar serves as the ants’ main food source and has coevolved with the ants for millions of years (12, 13, 15, 16). Ants forage for materials to feed to their cultivar fungus (e.g., grass, leaves, dried plant material, and/or insect frass), which the fungus degrades and converts to fungal biomass, including hyphal swellings called gonglydia that the ants eat as their primary food source (20, 21). *Pseudonocardia*, an Actinobacterium, grows on the cuticle of many fungus-growing ants, producing antimicrobials to protect the fungus garden from pathogens (22, 23). Other cuticular microbes on the ants have been identified but their function is unknown (23, 24). Together these interactions comprise a multipartite symbiosis composed of the fungus-growing ant, the *Leucoagaricus* fungus, and the protective symbiont *Pseudonocardia*.

Several studies have preliminarily investigated the fungus-growing ant microbiome, primarily focusing on leaf-cutting ants (25–30). These studies show that *Acromyrmex* and *Atta* leaf-cutting ants host low diversity microbiomes that include *Wolbachia, Solirubrobacter, Enterobacter, Pseudomonas*, and members of the orders Entomoplasmatales and Rhizobiales. Rhizobiales bacteria contain *nif* genes that were hypothesized to cycle nitrogen for the ants and their cultivar fungus (29), as did *Methylobacterium, Ralstonia* and *Pseudomonas* strains (30). *Wolbachia* is maternally transmitted in *Acromyrmex* and *Atta* but lacks a known function.

Entomoplasmatales bacteria belong to the genera *Mesoplasma* and *Spiroplasma* are thought to provide nutrients to the ants (28, 31). Most of these studies pooled multiple worker ants together before DNA extraction, leaving how ant microbiomes differ between individuals within a colony and between castes largely unknown.

Here, we characterize the microbiome of the fungus-growing ant *Trachymyrmex septentrionalis*. Phylogenetically, the higher agriculture ant genus *Trachymyrmex* holds a transitional position adjacent to leaf-cutting ants [17]. Typical *T. septentrionalis* colonies have approximately 1,000 workers in addition to a single queen, and during the summer months they also produce reproductive female and male alates that remain in the colony before mating as a mass swarm (16). *T. septentrionalis* colonies are found in pine barren and sandy soil ecosystems throughout the eastern USA and northern Mexico (20, 32, 33), and vary genetically across the United States, forming four different clades (33). Ishak et al (34) sampled *T. septentrionalis* ants from 25 colonies at a single location in Texas over one year and identified 19 bacterial genera that were conserved in *T. septentrionalis* workers and alates (both males and females). They also identified differences in ant microbiomes between castes, albeit including only a few samples.

However, the samples collected from each colony were not differentiated from each other, making it impossible to study intra-colony variation in microbiome composition. Whether the 19 common bacteria found in Texas *T. septentrionalis* and form a conserved microbiome that is found in other geographic regions or castes is also unknown.

In contrast to this previous work, we sampled individual ants collected across a broad geographic gradient to characterize the conserved microbiome of field-collected *T. septentrionalis* worker ants. We also more deeply sampled individual ant microbiomes within two colonies and found that ant gut microbiome composition differs between colonies, between ant castes, and following adaptation of colonies to the laboratory environment. The variable presence of *Mesoplasma* and *Spiroplasma* symbionts in different ant colonies and castes was a major driver of these microbiome differences. Our study provides an example of how individual colonies, castes, and ecological differences determine the composition of social insect microbiomes.

## METHODS

### Sample collection

*Trachymyrmex septentrionalis* colonies were collected in New Jersey, New York, North Carolina, Florida, Georgia, and Louisiana during 2014–2018 (Suppl. File 1; Suppl. Fig. S1). Permits for *T. septentrionalis* collection include: the New Jersey Department of Environmental Protection Division of Parks and Forestry State Park Service unnumbered letters of authorization; New York State Department of Environmental Conservation License to Collect or Possess: Scientific #915; Permit for Research in Suffolk County; North Carolina Division of Parks and Recreation Scientific Collection and Research Permit 2015_0030; Florida Department of Agriculture and Consumer Services unnumbered letters of authorization; Georgia Department of Natural Resources State Parks and Historic Sites Scientific Research and Collection Permit 032015; and Louisiana Department of Wildlife and Fisheries Permit WL-Research2016-10, in addition to permits from the United States Department of Agriculture for the transportation of ants to the University of Connecticut (USDA; P526P-14-00684). Freshly collected ants (hereafter: “field-collected ants”) were stored in DESS (20% dimethyl sulphoxide, 250 mM disodium EDTA, and saturated sodium chloride) (35) on dry ice immediately after colony collection in the field and then stored at -80 °C in the lab until sampling for DNA extraction. The remainder of each colony (including both fungus gardens and the remaining ants) was brought back to the University of Connecticut, kept in a USDA-approved and temperature-controlled room, and maintained following Sosa-Calvo et al (36). Colonies were placed in a 6-3/4” long x 4 13/16” width x 2-3/8″ height box that was lined with plaster of Paris, watered biweekly to maintain humidity, and provided with sterile cornmeal *ad libitum* as food for the fungus garden. Ants sampled from these colonies will be referred to as “lab-maintained”.

We generated two major datasets for this study. The first, which we will refer to as the “multi-state” dataset, included field-collected workers from all of the states mentioned above. This dataset was designed to test if ant microbiomes varied throughout the ants’ geographic range. The second, which we will refer to as the “two-colony” dataset, was made up of multiple ants sampled from Colony JKH000270, collected from Friendship NJ in July 2017, and Colony JKH000307, collected from Quaker Bridge NJ in July 2018. We sampled all available castes and life stages, including workers, pupae, larvae, male alates, and female alates, to test if microbiomes differed between castes and following lab adaptation. The colonies selected for the “two-colony” dataset were selected based on the availability of matching field and lab samples. Colony JKH000270 lab-maintained ants were sampled after a year and 4 months (some male alates were sampled earlier) and Colony JKH000307 lab-maintained ants were sampled after 4 months. Worker ants collected from the lab-maintained colonies were sampled immediately before DNA extraction without storage in DESS. Although lab-maintained samples were not stored in DESS, our previous research demonstrating that DESS effectively preserves the fungus garden microbiome suggests that this is to be unlikely a significant source of bias (35). Two confirmatory datasets were also used, with one including pupae and another dissected ant guts and whole ants sampled from the same colony (Suppl. Fig. S2 and S8).

### Sample preparation & DNA extraction

All ants were surface-cleaned using a 10 second submersion in 70% ethanol followed by a 10 second submersion in phosphate-buffered saline (PBS), repeated three times (37). Some pupae were ethanol-washed and others were not to determine if the Tenericute symbionts were located on their exoskeleton or internally. DNA was extracted from these surface-cleaned ants using the bead-beating and chloroform-isopropanol protocol described previously (35, 38). Our preliminary data showed minimal differences between microbiomes generated using whole surface-cleaned ants and ant guts, and so we used whole, surface-cleaned ants throughout this study for simplicity (Suppl. Fig. S2) while recognizing that actinobacteria in our dataset (predominantly *Phycicoccus* and *Pseudonocardia*) may be present due to incomplete surface cleaning. Negative controls containing only the DNA extraction reagents were processed alongside each batch of ant samples. The DNA concentration of each extract and negative control was determined using the Qubit dsDNA high-sensitivity assay protocol and a Qubit 3.0 fluorimeter (Invitrogen, Carlsbad CA).

### PCR screening & community amplicon sequencing

DNA samples were PCR amplified using primers 515F and 806R to amplify the 16S rRNA gene V4 region (39). Ten nanograms of template DNA was added to 5 µl Green GoTaq Reaction Mix Buffer (Promega, Madison, WI), 1.25 units GoTaq DNA Polymerase (Promega, Madison, WI), 10 µmol of each primer, and 300 ng bovine serum albumin (BSA; New England BioLabs Inc. Ipswitch, MA), to which nuclease-free H_2_O was added to reach a volume of 25 µl. Thermocycler conditions (BioRad, Hercules, CA) were: 3 min at 95°C, 30 cycles of 30 sec at 95°C, 30 sec at 50°C, and 60 sec at 72°C, followed by a 5 min cycle at 72°C and then an indefinite hold at 4°C. Gel electrophoresis confirmed the expected band size of 300–350 bp.

Samples that produced PCR products of the expected size and all negative controls were prepared for community amplicon sequencing of the 16S rRNA gene V4 region using an Illumina MiSeq at the University of Connecticut Microbial Analysis, Resources, and Services facility. Approximately 30 ng of DNA from each sample was added to a 96-well plate containing 10 µmol each of Illumina-barcoded versions of primers 515F and 806R, 5 µl AccuPrime buffer (Invitrogen, Carlsbad, CA), 50 mM MgSO_4_ (Invitrogen, Carlsbad, CA), 300 ng BSA (New England BioLabs Inc. Ipswitch, MA), a 1 µmol spike-in of both non-barcoded primers 515F and 806R, and 1 unit AccuPrime polymerase (Invitrogen, Carlsbad, CA), to which nuclease-free H_2_O was added to a volume of 50 µl. Reaction mixes were separated in a 384-well plate using an epMotion 5075 liquid handling robot (Eppendorf, Hamburg, Germany) forming three replicates (each with a volume of 16.7 µl). This plate was then transferred to a thermocycler (Eppendorf, Hamburg, Germany), which used the following conditions: 2 min at 95°C, 30 cycles of 15 sec at 95°C, 60 sec at 55°C, and 60 sec at 68°C, a final extension for 5 min at 68°C, and an indefinite hold at 4°C. Post-PCR, triplicate reactions were re-pooled using the epMotion and DNA concentrations were quantified using a QIAxcel Advanced capillary electrophoresis system (Qiagen, Hilden, Germany). Samples with concentrations > 0.5 ng/µl were pooled using equal DNA masses to create the final sequencing libraries. Libraries were then bead-cleaned using Mag-Bind RXNPure plus beads (OMEGA, Norcross, Georgia) in a 1:0.8 ratio of sequencing library to bead volume. Cleaned library pools were adjusted to a concentration of 1.1 ng/µl ± 0.1 ng/µl and their concentrations were confirmed using the Qubit dsDNA high-sensitivity assay on a Qubit 3.0 fluorimeter (Invitrogen, Carlsbad, California). Microbial community sequencing on an Illumina MiSeq using 2 × 250 bp libraries (Illumina, San Diego, California) was completed in 4 batches, each containing either the multi-state dataset (104 samples + 5 negative controls), the two-colony dataset (147 samples + 9 negative controls), the pupae dataset (20 samples + 1 negative control), or the whole ant/gut dissection dataset (10 samples + 1 negative control).

### Bioinformatic analyses

We analyzed all datasets individually. Reads were analyzed using R v3.5.3 (40) and the dada2 v1.11.1 (41) pipeline for amplicon sequence variants (ASVs) (https://benjjneb.github.io/dada2/tutorial.html, accessed: November 11, 2017). Read counts per sample were 5–1,952,121(multi-state dataset), 34–212,473 (two colony dataset), 15,495–92,408 (pupae dataset), and 17,565–132,790 (whole ant/gut dissection dataset; Suppl. File 1). Metadata files were imported into phyloseq v1.26.1 (42), creating a phyloseq R object that was used for subsequent analyses. Reads that were not classified as belonging to the kingdom Bacteria (i.e., those identified as Archaea or Eukaryote) using the SILVA database v128 (43, 44) were removed. The ASVs that matched to mitochondria were then removed separately because SILVA included them in the kingdom Bacteria. Samples were screened for contamination using the decontam v1.2.1 (45) prevalence protocol with a default threshold value of 0.1. No reads were flagged as contaminants for any of the datasets, resulting in 1,798, 1,353, 470, and 161 unique ASVs for the multi-state, two colony, pupae, and whole ant/gut dissection datasets, respectively (Suppl. File 1). Negative control samples were not considered further. All samples were rarefied to 20,000 reads for the multi-state dataset, 5,000 reads for the two-colony dataset, 10,000 reads for the pupae dataset, and 14,500 reads for the whole ant/gut dissection dataset, and read counts were converted to relative abundances. The final phyloseq objects contained 71, 110, 19, and 10 samples for the multi-state, two colony, pupae, and whole ant/gut dissection datasets, respectively.

Alpha diversity (Shannon and Simpson metrics) was measured using the phyloseq ‘plot_richness’ command and compared using the R stats ‘aov’ and ‘TukeyHSD’ commands. Beta diversity was measured using weighted Unifrac (WUF) and unweighted Unifrac (UUF) distance metrics. Weighted and unweighted Unifrac distances were calculated, ordinated, and viewed using the ‘distance’, ‘ordinate’, and ‘plot_ordinate’ phyloseq commands, respectively. Ellipses were added to the PCoA plots of each Unifrac distance using the ggplot2 (46) command ‘stat_ellipse’ with a default multivariate t-distribution. PERMANOVA tests were calculated using vegan v2.5-4 (47). Although weighted Unifrac and unweighted Unifrac distances were used for each test, to keep the text concise only one test is listed in the text and the complimentary values are presented in Suppl. Tables S2-S4. After exporting the multi-state phyloseq object into Microsoft Excel, we chose the first sample from each colony to be used in the Mantel tests, creating a new dataset containing 36 samples. Each sample and their corresponding GPS locations were uploaded to GeoMatrix (48) (Geographic Distance Matrix Generator v1.2.3) following the specified format. The GeoMatrix output was downloaded and imported into R. Full Mantel tests were completed using the GeoMatrix file and the WUF and UUF distance matrices for the 36 samples that included only one ant from each colony using the vegan command ‘mantel’. Partial Mantel tests were completed and Mantel correlograms created using the vegan ‘mantel.correlog’ and ‘plot’ commands, respectively. Kolmogorov-Smirnov and Wilcoxon rank sum tests were completed using the R commands “ks.test” and “wilcox.test”, respectively. If the p-value for the Kolmogorov-Smirnov test was > 0.05, then the Wilcoxon rank sum tests were used to compare the distributions of Unifrac distances; otherwise, a T-test was used (Suppl. Fig. S10B). A heatmap of the bacterial genera in the two-colony dataset whose read abundances were > 5% was created using the ‘heatmap.2’ command in gplots v3.0.3 (49). Heatmaps describing the median relative abundances of genera with a read abundance > 5% within workers and male alates from colony JKH000270 and JKH00307 were created using charticulator v2.0.4 (https://charticulator.com/app/index.html).

A phylogenetic tree was constructed to show relationships between the 16S rRNA sequences of the Tenericutes generated during this study and those that had been previously isolated from the leaf-cutting ants *Atta* and *Acromyrmex*. The most abundant *Spiroplasma* ASV and *Mesoplasma* ASV from the combined multi-state and two-colony datasets were aligned with reference Tenericute sequences generated from other ants and related *Mesoplasma* and *Spiroplasma* sequences, all downloaded from NCBI, using MUSCLE v3.8.31 (50) and then trimmed to the same alignment length. A phylogenetic tree, rooted using the *Bacillus subtilis* 16S rRNA gene sequence, was calculated using a GTRGAMMAI substitution model and 500 bootstrap replicates in RaxML v8.2.11 (51).

The commands used for all analyses are attached as Suppl. File 2. All raw sequencing reads are deposited in the NCBI database under BioProject PRJNA687229.

## RESULTS

### Ant microbiomes differ between states

We characterized *T. septentrionalis* worker ant microbiomes in field-collected ants from New York, down the U.S. East Coast into Florida, and westward into Louisiana (Suppl. Fig. S1). *T. septentrionalis* worker ant microbiomes were dominated by bacteria belonging to the phyla Actinobacteria, Firmicutes, Proteobacteria, and Tenericutes (Fig. 1A; Suppl. Fig. S3). Alpha diversity was low for these *T. septentrionalis* microbiomes and did not differ between states or between ants sampled from the same colony (Suppl. Fig. S4A & B; ANOVA: State p=0.492, Colony p=0.335). Eighteen genera were present with an abundance of ≥5% in at least one sample (Fig. 1A & B). The most prevalent genera in the dataset, making up the conserved *T. septentrionalis* worker ant microbiome, were *Thermomonas, Escherichia, Lautropia, Solirubrobacter, Pseudonocardia, Aeromicrobium*, and *Nocardioides* (Fig. 1A). These bacteria had a prevalence of at least 75% throughout the dataset and co-occurred frequently. *Phycicoccus, Mesoplasma, Spiroplasma, Acinetobacter*, and *Wolbachia* were less prevalent in this dataset (50– 75%; Fig. 1A). The relative abundances of all genera varied greatly between samples (Fig. 1B). *Pseudonocardia, Aeromicrobium, Nocardioides*, and *Solirubrobacter* each had high prevalence but low median relative abundances < 25% in most samples (Fig. 1B). *Thermomonas* and *Lautropia* had high median relative abundances that varied greatly between samples. When present, *Mesoplasma* and *Spiroplasma* were often highly abundant (> 50% of all microbes in a sample) and did not co-occur with one another in any sample. Many ant colonies sampled from New York and North Carolina lacked *Mesoplasma* or *Spiroplasma* (Suppl. Fig. S3). *Amycolatopsis, Bacillus*, and *Weissella* were also highly abundant in 1–2 samples (Fig. 1B).

**Figure 1).**
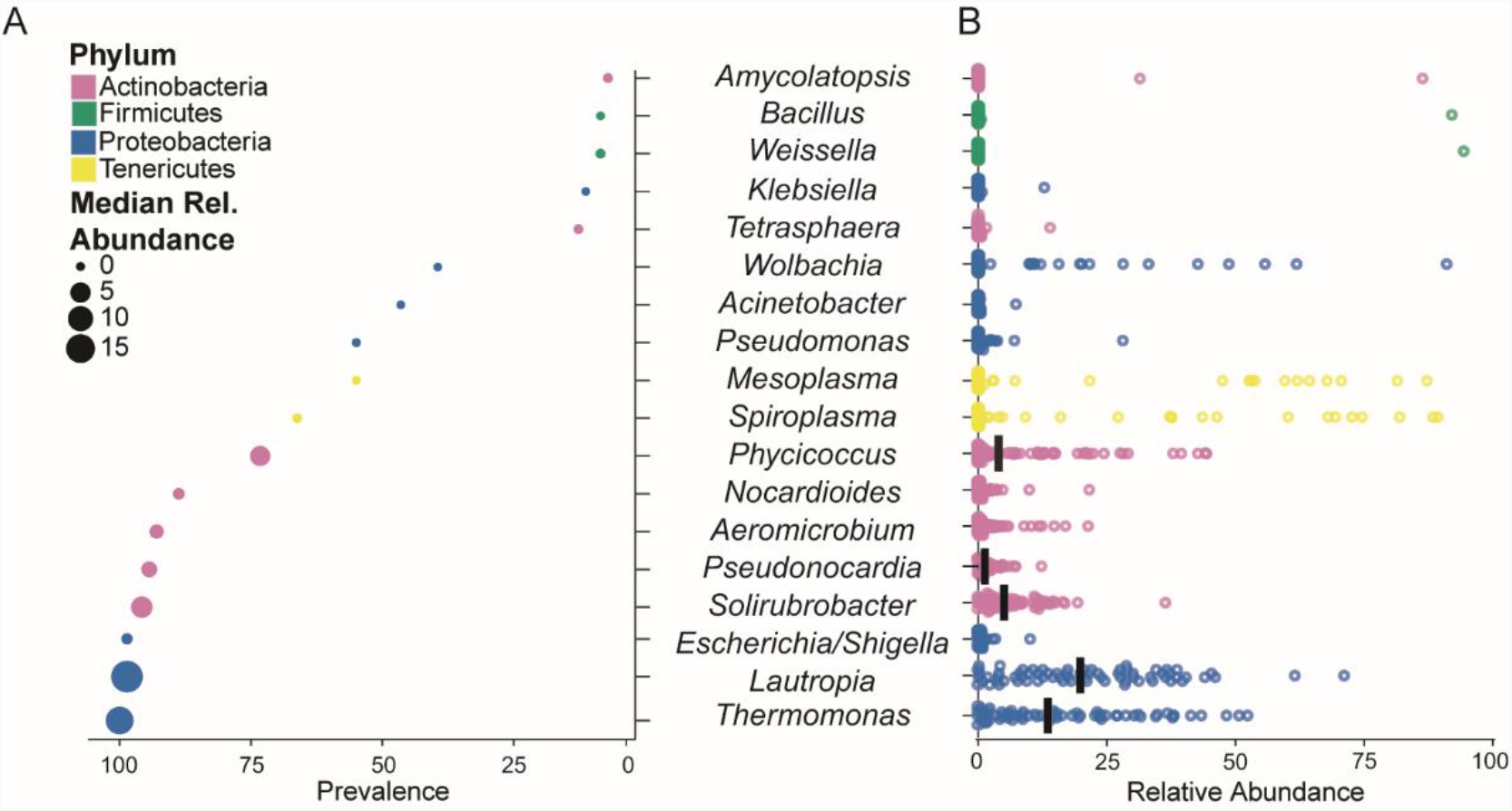
The *T. septentrionalis* worker ant conserved microbiome. A) Prevalence of bacterial genera in the multi-state dataset with an abundance ≥ 5% in at least one ant microbiome. Genera are ordered on the y-axis from most to least prevalent. Median relative abundance is represented by circle size and phylum is represented by color. B) Relative abundance of the bacterial genera ≥ 5% in at least one ant sample in the multi-state dataset, with the abundance in each sample represented by a single point for each genus. Bars indicate medians.

Multiple ants were collected from most colonies in the multi-state dataset, and the colony from which ants were collected explained more variation in microbiome composition than did the state of collection (WUF Colony PERMANOVA: R^2^= 0.524, p= 0.003; WUF State PERMANOVA R^2^= 0.135, p= 0.003; Suppl. Fig. S5B). These trends remained after excluding samples from New York and North Carolina, which lacked Tenericutes (Suppl. Fig. S5B). Although ant microbiomes from the multi-state dataset clustered weakly by state (Fig. 2; Suppl. Fig. S5A), weighted Unifrac distances between ant microbiomes did not correlate with the geographic distances between collection locations, although unweighted Unifrac distances did, albeit weakly (UUF Mantel R=0.143, p=0.019, Fig. 2B; WUF Mantel R= 0.034, p=0.778, Suppl. Fig. S5C & D). In the corresponding partial Mantel tests, distances between colonies from the smallest distance class (< 100km) were positively and significantly correlated to UUF distances between the ant microbiomes from these colonies (Fig. 2B; Suppl. Fig. S5D). Distance from classes 2 (∼250 km) and 5 (∼850 km) negatively and significantly correlated with UUF distances in ant microbiome composition, but all other correlations were non-significant (Fig. 2B; Suppl. Fig. 5C & D). Overall, these analyses show that microbiomes from colonies collected within 100 km from each other were somewhat similar to each other but that microbiomes otherwise varied in composition between colonies.

**Figure 2).**
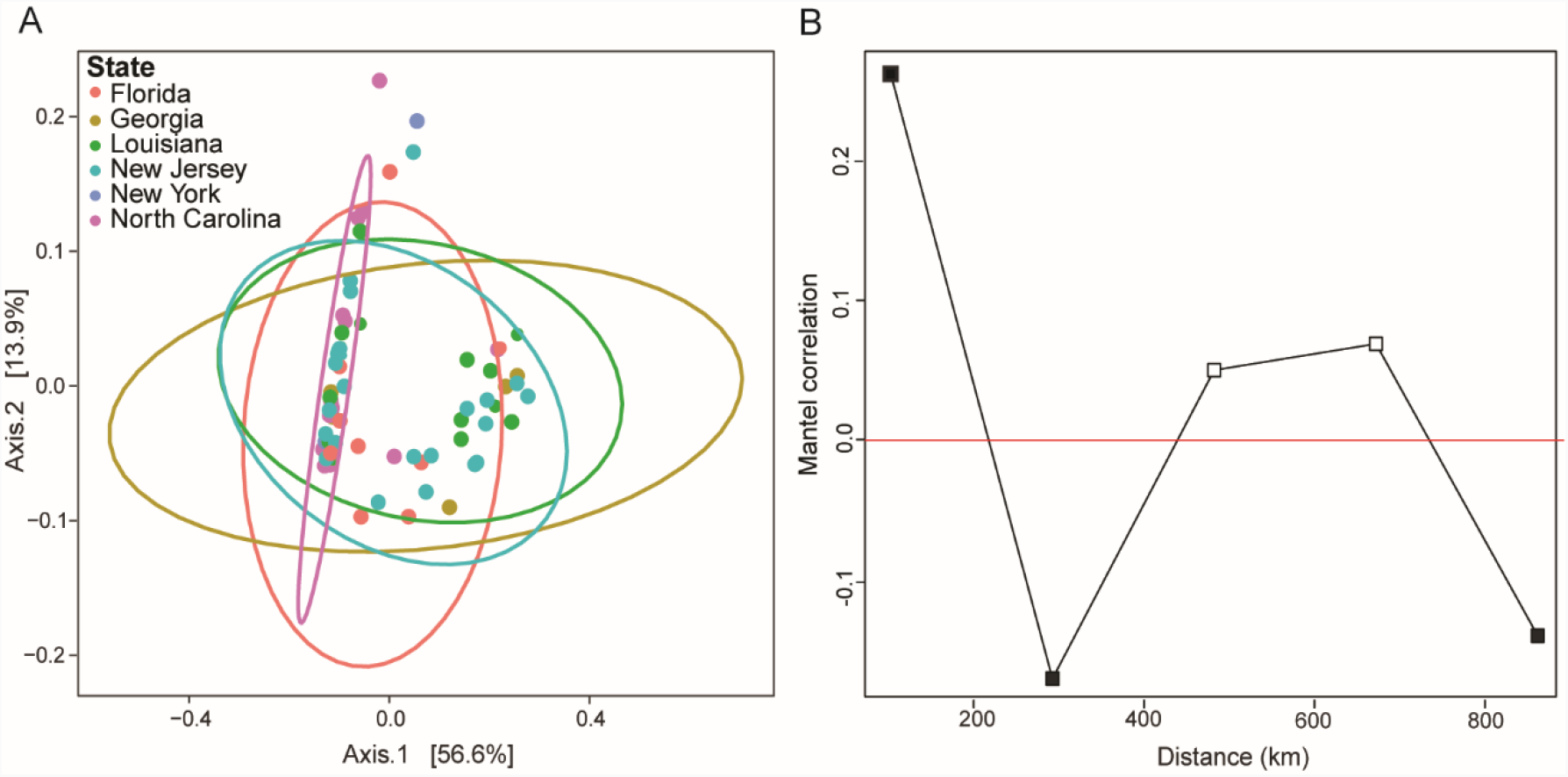
Biogeography of the *T. septentrionalis* worker microbiome. A) PCoA of Weighted Unifrac distances between ant microbiomes in the multi-state dataset. Samples are colored by state and grouped using a multivariate t-distribution. n=71. B) Partial Mantel correlation between Unweighted Unifrac distances and ant sampling locations in the multi-state dataset, excluding comparisons between ants sampled from the same colony. The x-axis indicates the distance class index in kilometers, and the y-axis indicates the Mantel correlation R statistic. Filled and unfilled squares indicate significant (p < 0.05) and nonsignificant (p > 0.05) p values, respectively, and points above and below the red line indicate positive and negative correlations, respectively.

### Ant microbiomes differ between castes and due to lab adaptations

In our multi-state dataset, we found that ant microbiomes were more conserved within a colony than between colonies. However, the multi-state dataset only included 1–3 field-collected worker ants to represent each colony. To overcome this limitation, we more deeply sampled individual ants from two colonies collected in Wharton State Forest, New Jersey (Suppl. File 1). The field-collected workers in this two-colony dataset had low alpha diversities, similar to those in the multi-state dataset (Suppl. Fig. S6A). We also found that the same phyla were conserved in the two-colony dataset as in the multi-state dataset. Field-collected worker ants were colonized by Proteobacteria, Tenericutes, and Actinobacteria, with some ants from colony JKH000270 also colonized with Bacteroidetes (Fig. 3A). Most conserved worker ant microbiome genera from the multi-state dataset (*Thermomonas, Lautropia, Solirubrobacter, Pseudonocardia, Aeromicrobium*, and *Nocardioides*) were also present in field-collected worker ants in the two-colony dataset (Suppl. Fig. S7).

**Figure 3).**
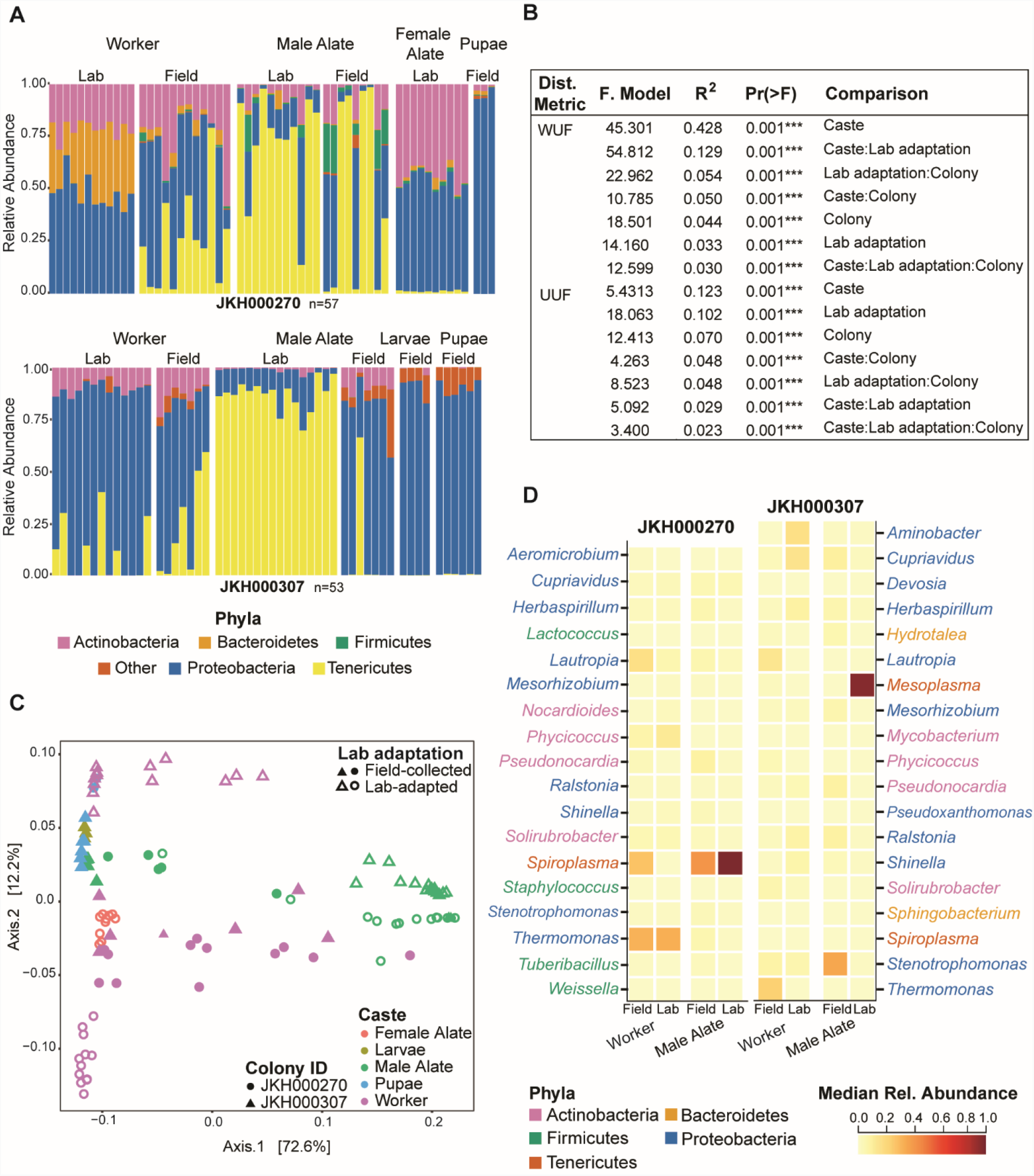
Colony, caste, and lab-adaptation all determine ant microbiome composition. A) Bar plot of the most abundant phyla in microbiomes from colonies JKH000270 (top) and JKH000307 (bottom). Single bars represent individual ant microbiomes, and sample types are grouped along the x-axis. The y-axis indicates relative abundance. Phyla are differentiated by color, and “other” represents phyla present at < 15% relative abundance. B) PERMANOVA analyses testing correlations between ant microbiome composition in the two-colony dataset and ant caste, lab adaptation status, and their colony of origin. n= 110. C) PCoA of Weighted Unifrac distances between ant microbiomes in the two-colony dataset. Colors indicate ant caste, shapes indicate colony ID, and solid and open shapes indicate field-collected and lab-maintained samples, respectively. D) Heatmap of genera with an abundance ≥ 5% in worker ants and male alates from colony JKH000270 (left) and colony JKH000307 (right). Genera are listed on the y-axis and colors indicate their phylum.

Although the limited number of colonies in the two-colony dataset is not ideally powered to compare ant microbiomes from different colonies in a general way (in contrast to the multi-state dataset), it instead prioritizes sampling of ants from different developmental stages and castes (pupae, larvae, male alates, and female alates) and ants collected before and after lab adaptation (field-collected or lab-maintained). Alpha diversity of lab-maintained male alates was often significantly lower compared to that of other sample types, which otherwise differed minimally from each other (Suppl. Fig. S6B). Like the workers, all castes were colonized by Actinobacteria, Proteobacteria, and Tenericutes, with Bacteroidetes only abundant in colony JKH000270, along with Firmicutes in some male alates (Fig. 3A). Compared to the worker microbiomes, Tenericutes were particularly abundant in male alate microbiomes, Proteobacteria in pupae and larvae, and Actinobacteria and Proteobacteria in female alates (Fig. 3A). There were extremely few *Mesoplasma* or *Spiroplasma* in pupae and larvae from the two-colony dataset. However, *Mesoplasma* and *Spiroplasma* were abundant in the separate pupae dataset generated from other *T. septentrionalis* colonies and used for confirmation (Suppl. Fig. S8A).

In the two-colony dataset, ant microbiome composition differed between all tested categories, and although caste explains the most variation it depends on the colony and lab adaptation (Fig. 3B & C). Caste accounted for the most microbiome variation using both WUF and UUF distances (Fig. 3B). These differences were colony-, caste-, and adaptation-specific because all interaction terms accounted for substantial microbiome variation using both WUF and UUF distances. Microbiomes from colonies JKH000270 and JKH000307 clustered separately along axis 2 in the WUF and UUF PCoAs but overlap in the center (Fig. 3C; Suppl. Fig. S9). This second axis accounts for a smaller amount of microbiome variation than axis 1, which separates the microbiomes into different castes, lab adaptation status, and combinations of these categories in both the WUF and UUF PCoAs.

Although each caste had a unique microbiome, we could only directly compare changes resulting from lab adaptation for workers and male alates, which were the only castes that we sampled from both field and lab settings (Fig. 3D). In colony JKH000270, field-collected and lab-maintained worker and male alate microbiomes differed principally in their abundances of *Spiroplasma* (Fig. 3D). In colony JKH000307, field-collected and lab-maintained worker microbiomes differed in their abundances of *Thermomonas, Lautropia, Aminobacter*, and *Cupriavidus*, and lab-maintained, but not field-collected, male alate microbiomes were dominated by *Mesoplasma* (Fig. 3D). Lab-maintained microbiomes were much less variable than the field-collected microbiomes, except for colony JKH000307 worker ants, when measured using UUF distances (Suppl. Fig. S10A & B).

In nearly all instances, microbiomes varied more between each category (colonies, caste, and lab adaption) than within the same category (Suppl. Fig. S10C). Microbiomes from each colony were grouped by caste, and then worker and male alate samples were further grouped into lab-maintained and field-collected categories to remove lab adaptation as a potentially confounding variable (Suppl. Fig. S10C). Typically, median WUF and UUF distances were greater between categories (caste, colony, lab adaptation) than within them. However, this was not true for UUF JKH000270 field-collected male alates, JKH000270 female alates, and JKH000307 larvae (Suppl. Fig. S10C). Overall, these comparisons highlight that ant microbiomes are more similar within our sampling categories (colony, caste, lab adaptation) than between them.

### *Spiroplasma* and *Mesoplasma* are *T. septentrionalis* symbionts

*Mesoplasma* and *Spiroplasma* were the two dominant genera in our microbiome samples, accounting for 12% and 14% of all the reads in our datasets, respectively. These bacteria, classified within the phylum Tenericutes, are well-known symbionts of insects and plants (52–57). Although their function remains to be experimentally tested, *Spiroplasma* and *Mesoplasma* have been found in other fungus-growing ants (26–29, 31, 34). The most abundant *Mesoplasma* ASV in our dataset, comprising 97.7% of all *Mesoplasma* reads, was highly similar to the 16S rRNA of EntAcro1 and an uncultured Entomoplasmataceae 16S rRNA sequence found in army ants (Fig. 4) (58). EntAcro1 is a metagenome-assembled genome (MAG) that was assembled from *Acromyrmex* fungus-growing ants (31), and related 16S rRNA sequences have also been found in *Atta colombica* and *At. sexdens, Mycetomoellerius zeteki* (previously *Trachymyrmex zeteki*), *Paratrachymyrmex cornetzi* (previously *T. cornetzi*) and *Cyphomyrmex rimosus* fungus-growing ants. The most abundant *Spiroplasma* ASV in our dataset, comprising 97.3% of all *Spiroplasma* reads, was highly similar to the EntAcro10 MAG assembled from *Acromyrmex* ants (31) and for which related 16S rRNA sequences were found in *Atta, Acromyrmex*, and *Mycetomoellerius* (27, 29) fungus-growing ants, along with other 16S rRNA sequences that were annotated as uncultured Entomoplasmatales from two other fungus-growing ants, *Sericomyrmex amabilis* and *M. jamaicensis* (Fig. 4) (26, 59).

**Figure 4).**
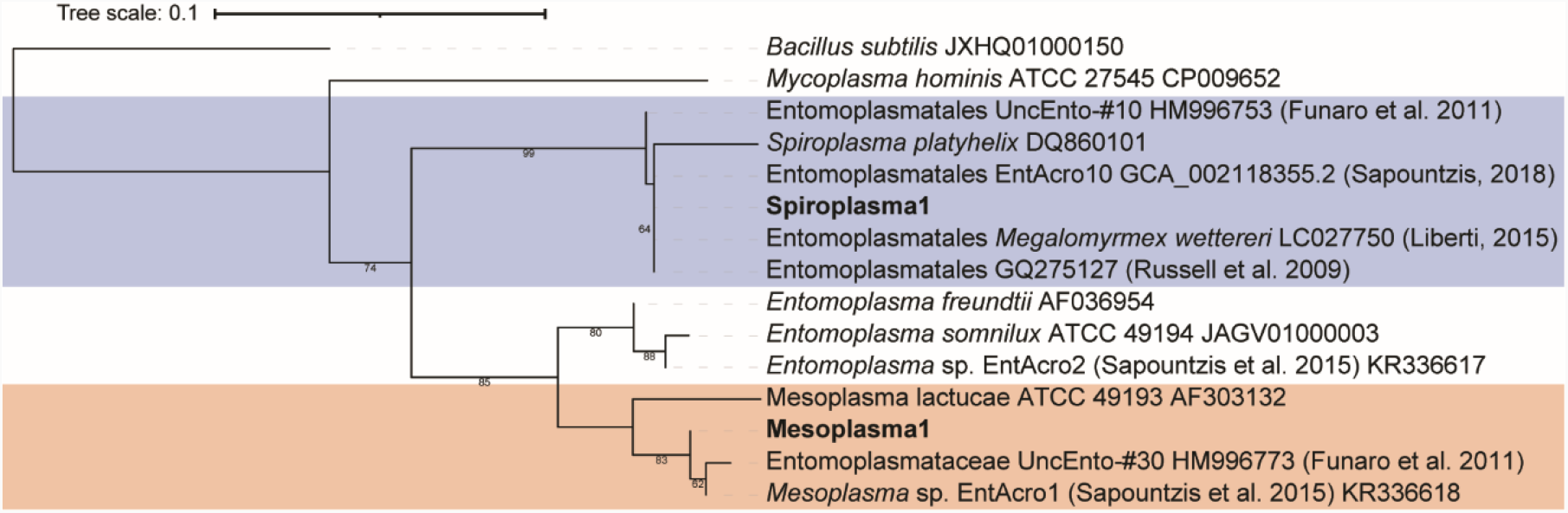
Maximum-likelihood phylogenetic tree of the dominant Tenericute 16 rRNA gene ASVs from the combined multi-state and two-colony datasets (in bold text) and reference 16S rRNA gene sequences sequenced from other fungus-growing ants and related Tenericutes strains. The tree was rooted using *Bacillus subtilis* and *Mycoplasma hominis* and bootstrap percentages > 60 are shown.

The presence of *Mesoplasma* and *Spiroplasma* varied between different ant colonies, between ants from the same colony, and between ants from different castes, both before and after lab adaptation. In our multi-state dataset, colonies from New York and North Carolina had lower Tenericute abundances compared to the other states (Fig. 5A). *Spiroplasma* was abundant in field-collected worker ants and both field-collected and lab-maintained male alates from colony JKH000270. *Mesoplasma* was abundant in field-collected and lab-maintained workers and lab-maintained male alates from colony JKH000307. In both the multi-state and two-colony datasets, only one of *Spiroplasma* or *Mesoplasma* was present per colony, although not all ants from that colony were colonized by a Tenericute (Fig. 5B & C). An exception to this trend was in Colony JKH000307, in which *Mesoplasma* was abundant but *Spiroplasma* colonized two lab-maintained male alates and two field-collected workers. Ants colonized by *Spiroplasma* were rarely found within a colony mainly colonized by *Mesoplasma*, and vice versa (Fig. 5B & C), and no ants in our multi-state or two-colony datasets were colonized by both *Spiroplasma* and *Mesoplasma*.

**Figure 5).**
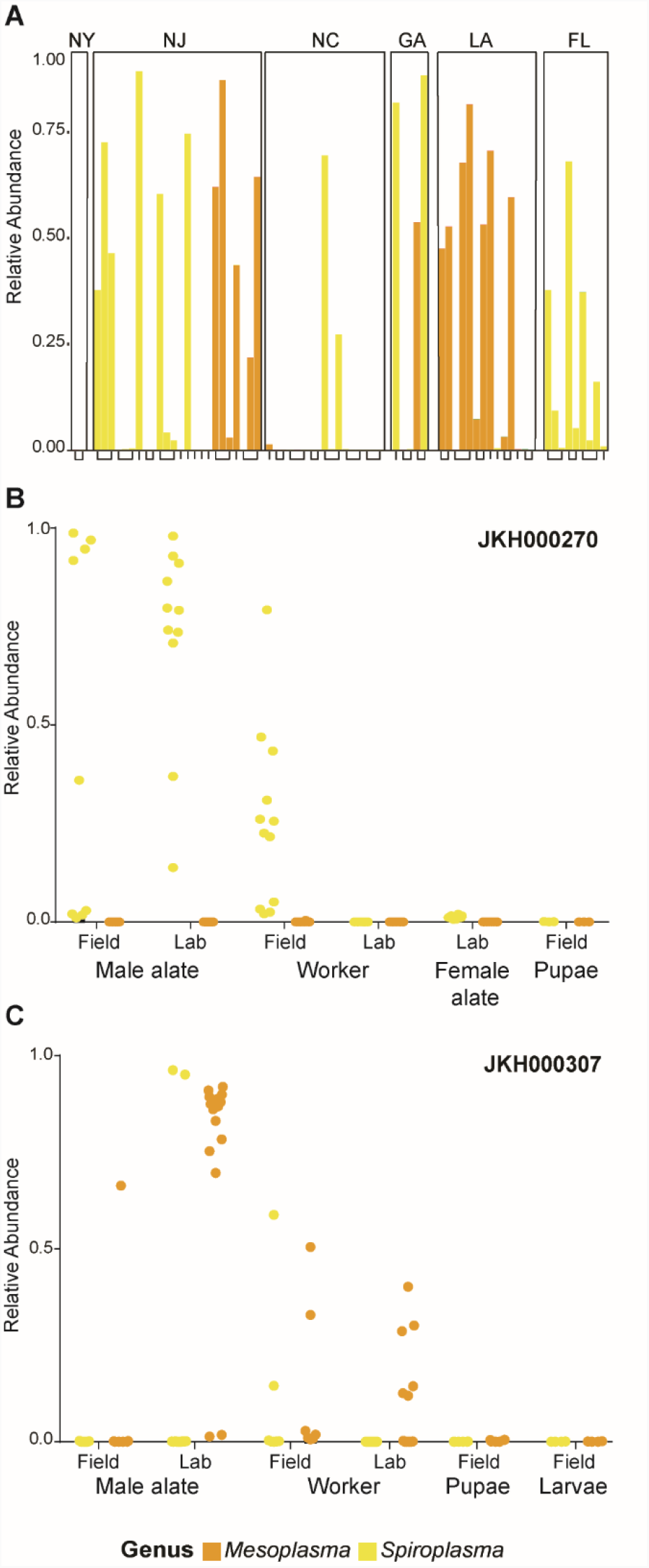
*Spiroplasma* (yellow) and *Mesoplasma* (orange) are abundant in *T. septentrionalis* ants. A) Single bars represent the relative abundance of reads from the genus *Spiroplasma* or *Mesoplasma* in ants from the multi-state dataset, grouped by the state of collection. X-axis brackets group worker ants from the same colony. n=71. B & C) The relative abundance of the genus *Spiroplasma* or *Mesoplasma* in the two-colony dataset, with each dot representing relative abundance in a single ant. All ants in this dataset were colonized exclusively by either *Spiroplasma* or *Mesoplasma*.

However, in a confirmatory dataset that sampled pupae from two additional ant colonies JKH000125 and JKH000136, and we found that pupae from colony JKH000136 were colonized exclusively by large amounts of *Mesoplasma* and that pupae from colony JKH000125 pupae were colonized primarily by *Spiroplasma* but also by small amounts of *Mesoplasma* (Suppl. Fig. S8C). This is the only time we have observed ants colonized by both *Mesoplasma* and *Spiroplasma*.

## DISCUSSION

Our microbiome sequencing of individual *T. septentrionalis* ants shows that caste, lab adaptation (field-collected or lab-maintained), and the colony that an ant belongs to all influence the structure of the ant microbiome. Our multi-state dataset includes field-collected workers ants from 6 states representing 36 colonies, allowing us to describe a conserved microbiome of wide-ranging worker ants that contains *Thermomonas, Escherichia, Lautropia, Solirubrobacter, Pseudonocardia, Aeromicrobium*, and *Nocardioides* (Fig. 1A), all of which were prevalent in > 75% of the ants in this dataset. This conserved microbiome was largely confirmed in the field-collected worker ant microbiome in our two-colony dataset. *Wolbachia* was also found in several samples from our multi-state dataset, similar to what has been documented in *Acromyrmex*, where it is thought to be transmitted maternally (28). *Mesoplasma* and *Spiroplasma* were both found throughout our dataset (Fig. 5) and are also abundant in other Attine ant genera (27, 28). Our results also agree with those of Ishak et al. (34), who identified 19 bacterial genera that were common in several *T. septentrionalis* colonies sampled from a single location in Texas. Of these 19 genera, 9 are among the most abundant 18 genera in our multi-state dataset: *Aeromicrobium, Nocardioides, Phycicoccus, Pseudonocardia, Solirubrobacter, Bacillus, Pseudomonas, Mesoplasma*, and *Spiroplasma*. The larger number of taxa found by Ishak et al. (34) is consistent with our finding that the *T. septentrionalis* ant microbiome has some biogeographic conservation, albeit limited. Colonies that are located closer together had the most similar microbiome compositions, meaning that a list of common taxa sampled from a single collection location would be larger than a conserved microbiome defined using samples collected from many different locations.

In our multi-state dataset, in which 1-3 ants were sampled per colony, the microbiomes from ants sampled from the same colony were more similar than ants sampled from different colonies. To further study the variation between the microbiomes of individual ants from a single colony, our two-colony dataset included microbiomes from ants more deeply sampled from two colonies, including ants from different castes and from both before and after lab adaptation. Again, we found that ant microbiomes within colonies were more similar than those between colonies. Despite differences in their modes of microbiome transmission and selection, this bolsters a common natural pattern, e.g., where nestmates of birds (60) and mammals (61), and colonies of insects such as honey bees (9, 11) and ants (62) all have similar microbiomes. These differences that we observed between colonies could be due to local ecological factors such as regional differences in the plants, insect frass, and the microbes associated with these substrates that will cause the ant microbiome to vary. The soil microbiome that surrounds the fungus garden chambers is highly conserved and overlaps minimally with the fungus garden microbiome (Lee and Klassen, unpublished data), suggesting that input from soil microbes is unlikely to drive significant differences in ant microbiomes. This is supported by our samples collected following adaptation of colonies to our lab, where environmental conditions are constant and colonies are fed only sterile cornmeal. Lab-maintained ant microbiomes are much less variable than those sampled in the field (Suppl. Fig. S10B), highlighting the importance of natural foraging and environmental variation for maintaining microbiome diversity. Another possible reason why microbiomes might differ between colonies is the strong founder effects that likely occur during colony foundation. Before leaving to mate, a female alate will place a piece of the fungus garden in her infra-buccal pocket (13). After mating, this new queen will found her own colony using that piece of fungus garden, tending it and producing all offspring in that colony. The microbiome possessed by the queen will be passed to these workers who will thereafter pass it on to subsequent generations of workers, thus creating a bottleneck in microbiome composition caused by the limited microbial diversity possessed by the queen during colony foundation.

Within a colony, microbiome composition depended on ant castes, which determined the greatest amount of variation between microbiome composition in the two-colony dataset (Fig. 3B). Gut microbiomes often differ between social insect castes, likely due to the specialized functional role played by each caste (2). Each insect in a colony will shed their gut lining during pupation, thereby also shedding their gut microbiome and requiring the gut microbiome to then reassemble. The microbiomes of reproductive castes often differ from those of the worker castes in termites (8), honey bees (9), and other species of ants (62). In *T. septentrionalis* colonies, workers forage for plant material and caterpillar frass that have their own microbiomes, and these bacteria might be transferred to the workers and to the fungus garden that they tend. Such microbes brought into the colony via foraging could then be seen transferred to all ant microbiomes in that colony. However, such transfer may be rare due to the ants grooming themselves, other ants, and the fungus garden to remove pathogens (63). In contrast, the role of male and female alates is to mate and, for females, to start a new colony. The different microbiomes possessed by these castes could therefore reflect their need to reproduce, which the workers do not experience. Female alates possess large amounts of lipids, proteins, and carbohydrates to provide energy when they start their colonies (64), and male alates must travel further than female alates to mate (65). The different microbiome composition possessed by male and female alates (Fig. 3C) could therefore reflect the differing energy and nutrient requirements of their different reproductive behaviors (64, 65). Although we show that *T. septentrionalis* microbiomes differ between castes, how vertical versus environmental transmission contributes to these differences remains unknown.

The presence and abundance of *Mesoplasma* and *Spiroplasma* differ between each colony and individual ant, explaining some of the differences in ant microbiome composition between ant colonies and castes. The most abundant *Mesoplasma* and *Spiroplasma* 16S rRNA genes detected in our dataset are very similar to those of other Tenericutes found in other fungus-growing ant genera, which suggests that these bacteria are a common member of the fungus-growing ant symbiosis. The functional role played by these Tenericutes is unknown. They are unlikely to be reproductive pathogens, based on their high abundances in worker ants, which are reproductive dead ends. *Acromyrmex* and *Atta* callows (i.e., newly emerged worker ants) were readily colonized by Tenericutes when cared for by other Tenericutes-containing worker ants, but not after 21 days in the absence of such ants (28). Thus, Tenericutes symbionts are likely transferred between ants via care from other workers. FISH imaging has also been used to localize Tenericutes to ant guts but not in reproductive organs. Because we only have characterized female alate microbiomes from a single colony, future studies should include a wider sampling of this caste. Tenericutes might also be acquired from an environmental source (e.g., foraged plants or caterpillar frass), or during interactions with other ant colonies (66).

Tenericute symbionts of fungus-growing ants may be nutritional symbionts. Metagenomic reconstructions of *Spiroplasma* and *Mesoplasma* symbionts from *Acromyrmex* ants suggested that these bacteria may decompose excess arginine, and for *Mesoplasma*, citrate (31). Arginine recycling would produce ammonium that could be provided to the fungus garden via ant fecal droplets, and the degradation of citrate, acquired from foraged fruit, leaves, and other plant material, could produce acetate for use by the ants. If these hypotheses were supported, it would suggest that Tenericutes provision nutrients for both the ants and the cultivar fungus. In the phylogeny of fungus growing ants, *Trachymyrmex* is the closest phylogenetic neighbor to the leaf-cutting ants (14, 67, 68). The fact that both the *Mesoplasma* and *Spiroplasma* found in this study are closely related to those found in *Atta* and *Acromyrmex*, unlike in more basal fungus-growing ant taxa, reflects this phylogenetic relationship. However, in our data Tenericute presence varied extensively between ant colonies and castes (Fig. 5), meaning that these symbionts must not be absolutely required for ant survival, especially given their complete absence in some ants, a hypothesis to be tested in future studies. Instead, such extreme variation in abundance between colonies and individual ants could imply that *Mesoplasma* or *Spiroplasma* are ant pathogens or commensals, at least in some ecological contexts. Interestingly, we did not find any co-infections of both *Mesoplasma* and *Spiroplasma* together in the same ant, except for in pupae from a single ant colony (Suppl. Fig. 8C). This might suggest a transmission bottleneck during microbiome assembly that allows for colonization of only one of these symbionts, or competition between *Spiroplasma* and *Mesoplasma* in these ants.

Our study shows how microbiome assembly differs between ant colonies, castes and as a result of lab adaptation. *T. septentrionalis* ant microbiomes did not vary extensively between sampling locations but did vary substantially between ant colonies and castes. Lab adaptation showed the importance of ant foraging or environmental variation for maintaining microbiome diversity, with lab-maintained microbiomes being less variable than field-collected ones as a reflection of their more stable environment. The presence of *Mesoplasma* and *Spiroplasma* cause some of these differences between colonies, castes, and as the result of lab-adaptation. Together, these data show how the composition of a social insect microbiome is the result of colony- and caste-specific factors.

## Acknowledgments

We would like to thank the state forest and park staff for their assistance with our fieldwork, the members of the Klassen Lab for their help collecting ant colonies and for their thoughtful feedback on this manuscript, and the UConn Microbial Analysis, Resources, and Services facility for microbiome sequencing. This research was funded by the U.S. National Science Foundation (IOS-1656475) and the University of Connecticut.

