## Supplemental File 2 for "*Trachymyrmex septentrionalis* ant microbiome assembly is unique to individual colonies and castes"

```
TRACHYMYRMEX WHOLE ANTS vs DISSECTED GUTS
#Load dada2, I used v1.11.1  and followed their published workflow
library(dada2)
path <- "~/Prelim Guts vs WA/"
fnFs <- sort(list.files(path, pattern="_R1_001.fastq", full.name = TRUE))
fnRs <- sort(list.files(path, pattern="_R2_001.fastq", full.name = TRUE))
sample.names <- sapply(strsplit(basename(fnFs), "_"), `[`, 1)
#View quality plots and I ended up adjusted "truncLen" based on this.
plotQualityProfile(fnFs[1:2])
plotQualityProfile(fnRs[1:2])
filt_path <- file.path(path, "filtered")
filtFs <- file.path(filt_path, paste0(sample.names, "_F_filt.fastq"))
filtRs <- file.path(filt_path, paste0(sample.names, "_R_filt.fastq"))
out <- filterAndTrim(fnFs, filtFs, fnRs, filtRs, truncLen = c(240,160), maxN=0, maxEE = c(2,5), truncQ=2, rm.phix =TRUE, compress=TRUE, multithread = TRUE)
errF <- learnErrors(filtFs, multithread = TRUE)
errR <- learnErrors(filtRs, multithread = TRUE)
#View Errors
plotErrors(errF, nominalQ=TRUE)
derepFs <- derepFastq(filtFs, verbose=TRUE)
derepRs <- derepFastq(filtRs, verbose=TRUE)
names(derepFs) <- sample.names
names(derepRs) <- sample.names
dadaFs <- dada(derepFs, err=errF, multithread=TRUE)
dadaRs <- dada(derepRs, err=errR, multithread=TRUE)
dadaFs[[1]]
mergers <- mergePairs(dadaFs, derepFs, dadaRs, derepRs, verbose = TRUE)
seqtab <- makeSequenceTable(mergers)
seqtab.nochim <- removeBimeraDenovo(seqtab, method="consensus", multithread=TRUE, verbose = TRUE)
sum(seqtab.nochim)/sum(seqtab)
getN <- function(x) sum(getUniques(x))
track <- cbind(out, sapply(dadaFs, getN), sapply(mergers, getN), rowSums(seqtab), rowSums(seqtab.nochim))
colnames(track) <- c("input", "filtered", "denoised", "merged", "tabled", "nonchim")
rownames(track) <- sample.names
#Add in taxa information using your choice of database.
taxa <- assignTaxonomy(seqtab.nochim, "Prelim Guts vs WA/silva_nr_v128_train_set.fa", multithread = TRUE)
taxa.print <- taxa
rownames(taxa.print) <- NULL
head(taxa.print)
#Output the taxa, taxa counts, and taxa asv sequences
write.csv(taxa.print, "AntGutWholeAnttaxa.csv")
write.csv(seqtab.nochim, "AntGutWholeAntASV.csv")

#Load other packages
library(phyloseq)
library(ggplot2)
library(plyr)
library(vegan)
#Read in Metadata. Includes the sample name, whether it is a sample or control, and other information that is needed. I included whether they were whole ants or dissected guts
meta <- read.csv("~/Prelim Guts vs WA/AGWA-metadata.csv",header = TRUE, row.names = 1)
ps <- phyloseq(otu_table(seqtab.nochim, taxa_are_rows = FALSE), sample_data(meta), tax_table(taxa))
#Check phyloseq object
ps
#Load decontam package
library(decontam)
sample_variables(ps)
df <- as.data.frame(sample_data(ps))
df$LibrarySize <- sample_sums(ps)
df <- df[order(df$LibrarySize),]
df$Index <- seq(nrow(df))
ggplot(data=df, aes(x=Index, y=LibrarySize, color=Sample_or_Control)) + geom_point()
sample_data(ps)$is.neg <- sample_data(ps)$Sample_or_Control == "Control Sample"
contamdf.prev <- isContaminant(ps, method="prevalence", neg="is.neg")

table(contamdf.prev$contaminant)

#To remove Eukyota, Archaea
ps1 <- subset_taxa(ps,Kingdom == "Bacteria")
#View it
get_taxa_unique(ps1, taxonomic.rank = "Kingdom")
#remove Blank sample
ps2 <- subset_samples(ps1 , sample_names(ps1) != "Blank")
#Remove anything not identified at Phylum level
ps2 <- subset_taxa(ps2, !is.na(Phylum))
sort(get_taxa_unique(ps2, taxonomic.rank = "Phylum"))
#Remove Mitochondria
ps3 <- subset_taxa(ps2, !Family == "Mitochondria")
sort(get_taxa_unique(ps3, taxonomic.rank = "Family"))
ps3

#Rarefy to 14500 reads
rngseed = 5
ps4 <- rarefy_even_depth(ps3, sample.size = 14500, rngseed = TRUE)
#`set.seed(TRUE)` was used to initialize repeatable random subsampling.
#Please record this for your records so others can reproduce.
#Try `set.seed(TRUE); .Random.seed` for the full vector
#9OTUs were removed because they are no longer
#present in any sample after random subsampling

#Put into relative abundance
psPhyla.rab <- transform_sample_counts(ps4, function(x) x/sum(x))
#Glom asvs of the same genus together
psPhyla.rab.glom <- tax_glom(psPhyla.rab, taxrank = "Phylum")
psPhyla.rab.glom.df <- psmelt(psPhyla.rab.glom)
#Plot phylum level barplot
psPhyla.rab.glom.df$Phylum <- as.character(psPhyla.rab.glom.df$Phylum)

psPhyla.barplot.phylum <- ggplot(psPhyla.rab.glom.df, aes(x = Sample, y = Abundance, fill = Phylum, scale_color_brewer(pallette = "Set3"))) + geom_bar(stat = "identity") + theme(axis.text.x = element_text(angle = 270, vjust = 0.5))
plot(psPhyla.barplot.phylum)
#Arrange into whole ant or dissected gut
psPhyla.barplot.phylum <- ggplot(psPhyla.rab.glom.df, aes(x = Sample, y = Abundance, fill = Phylum, scale_color_brewer(pallette = "Set3"))) + geom_bar(stat = "identity") + theme(axis.text.x = element_text(angle = 270, vjust = 0.5)) + facet_grid(~Type, scales = "free", space = "free") + theme(strip.text = element_text(angle = 315), strip.background = element_blank(), panel.background = element_blank())
plot(psPhyla.barplot.phylum)
#Top Phyla arranged by whole ant or dissected guts
max <- ddply(psPhyla.rab.glom.df, ~Phylum, function(x) c(max = max(x$Abundance)))
Other <- max[max$max <= 0.15,]$Phylum
psPhyla.rab.glom.df[psPhyla.rab.glom.df$Phylum %in% Other,]$Phylum <- "Other"
psPhyla.barplot.phylum <- ggplot(psPhyla.rab.glom.df, aes(x = Sample, y = Abundance, fill = Phylum, scale_color_brewer(pallette = "Set3"))) + geom_bar(stat = "identity") + theme(axis.text.x = element_text(angle = 270, vjust = 0.5)) + facet_grid(~Type, scales = "free", space = "free") + theme(strip.text = element_text(angle = 315), strip.background = element_blank(), panel.background = element_blank())
plot(psPhyla.barplot.phylum)


#Bray Curtis NMDS
psBray <- subset_samples(psPhyla.rab)
psBray_bcdist <- distance(psBray, method ="bray")
psBray_bcord <- ordinate(psBray, method= "NMDS", distance = "bray")
psBray_plotord <- plot_ordination(psBray, psBray_bcord, color = "Type")
plot(psBray_plotord)
#Permanova
psBray_df <- data.frame(sample_data(psBray))
View(psBray_df)
adonis(psBray_bcdist ~Type, data = psBray_df)

#PCOA
psV <- psPhyla.rab
psV.df <- t(data.frame(otu_table(psV)))
psV.df <- data.frame(psV.df)
psV.df$abundance <- rowSums(psV.df[,c(1:10)])
psV.df$sequence <- rownames(psV.df)
psV.uniques <- getUniques(psV.df)
uniquesToFasta(psV.uniques, "WAAG.fasta", ids = psV.df$sequence)
##After uniquesToFasta command, align seqs of the fasta file and then create phylogenetic tree. upload BestTree into read_tree command. For this publication I used CIPRES (available at https://www.phylo.org/portal2/login!input.action). Sequences were aligned using MAFFT on XSEDE (7.402), and phylogenetic tree made using RAxML-HPC BlackBox (8.2.12).
#Make Tree files
V.tree <- read_tree("~/Prelim Guts vs WA/RAxML_bestTree.result")
psV.tree <- phyloseq(otu_table(otu_table(psV)),sample_data(sample_data(psV)), tax_table(tax_table(psV)), phy_tree(V.tree))

#Unweighted unifrac
V.uuf.dist <- distance(psV.tree, method = "unifrac")
V.uuf.ord <- ordinate(psV.tree, method = "PCoA", distance = "unifrac")
V.uuf.plot <- plot_ordination(psV.tree, V.uuf.ord, color = "Type")
plot(V.uuf.plot)

#PERMANOVA
psV.tree.data <- data.frame(sample_data(psV.tree))
adonis(V.uuf.dist ~Type, psV.tree.data)

#Weighted unifrac
V.wuf.dist <- distance(psV.tree, method = "wunifrac")
V.wuf.ord <- ordinate(psV.tree, method = "PCoA", distance = "wunifrac")
V.wuf.plot <- plot_ordination(psV.tree, V.wuf.ord, color = "Type")
plot(V.wuf.plot)

#PERMANOVA
psV.tree.data <- data.frame(sample_data(psV.tree))
adonis(V.wuf.dist ~Type, psV.tree.data)

MULTI-STATE DATASET AND TWO-COLONY DATASET COMBINED
library(dada2)
path <- "~/MixedTrachyDataset/"
fnFs <- sort(list.files(path, pattern="_R1_001.fastq", full.name = TRUE))
fnRs <- sort(list.files(path, pattern="_R2_001.fastq", full.name = TRUE))
sample.names <- sapply(strsplit(basename(fnFs), "_"), `[`, 1)
#View quality plots and I ended up adjusted "truncLen" based on this.
plotQualityProfile(fnFs[1:2])
plotQualityProfile(fnRs[1:2])
filt_path <- file.path(path, "filtered")
filtFs <- file.path(filt_path, paste0(sample.names, "_F_filt.fastq"))
filtRs <- file.path(filt_path, paste0(sample.names, "_R_filt.fastq"))
out <- filterAndTrim(fnFs, filtFs, fnRs, filtRs, truncLen = c(240,160), maxN=0, maxEE = c(2,5), truncQ=2, rm.phix =TRUE, compress=TRUE, multithread = TRUE)
errF <- learnErrors(filtFs, multithread = TRUE)
errR <- learnErrors(filtRs, multithread = TRUE)
#View errors
plotErrors(errF, nominalQ=TRUE)
derepFs <- derepFastq(filtFs, verbose=TRUE)
derepRs <- derepFastq(filtRs, verbose=TRUE)
names(derepFs) <- sample.names
names(derepRs) <- sample.names
dadaFs <- dada(derepFs, err=errF, multithread=TRUE)
dadaRs <- dada(derepRs, err=errR, multithread=TRUE)
dadaFs[[1]]
mergers <- mergePairs(dadaFs, derepFs, dadaRs, derepRs, verbose = TRUE)
seqtab <- makeSequenceTable(mergers)
seqtab.nochim <- removeBimeraDenovo(seqtab, method="consensus", multithread=TRUE, verbose = TRUE)
sum(seqtab.nochim)/sum(seqtab)
getN <- function(x) sum(getUniques(x))
track <- cbind(out, sapply(dadaFs, getN), sapply(mergers, getN), rowSums(seqtab), rowSums(seqtab.nochim))
colnames(track) <- c("input", "filtered", "denoised", "merged", "tabled", "nonchim")
rownames(track) <- sample.names
#Add in taxa information using your choice of database.
taxa <- assignTaxonomy(seqtab.nochim, "~/MixedTrachyDataset/silva_nr_v128_train_set.fa", multithread = TRUE)
taxa.print <- taxa
rownames(taxa.print) <- NULL
head(taxa.print)
#Output the taxa, taxa counts, and taxa asv sequences
write.csv(taxa.print, "TaxaListMixedTrachy.csv")
write.csv(seqtab.nochim, "NameListMixedTrachy.csv")

#Load other packages
library(phyloseq)
library(ggplot2)
library(plyr)
library(vegan)
#Read in Metadata. Includes the sample name, whether it is a sample or control, and other information that is needed.
meta <- read.csv("~/MixedTrachyDataset/MixedTrachyMetaData.csv",header = TRUE, row.names = 1)
ps <- phyloseq(otu_table(seqtab.nochim, taxa_are_rows = FALSE), sample_data(meta), tax_table(taxa))

#To remove Eukyota, Archaea
ps1 <- subset_taxa(ps,Kingdom == "Bacteria")
#View it
get_taxa_unique(ps1, taxonomic.rank = "Kingdom")
#Remove anything not identified at Phylum level
ps2 <- subset_taxa(ps1, !is.na(Phylum))
sort(get_taxa_unique(ps2, taxonomic.rank = "Phylum"))
#Remove Mitochondria
ps3 <- subset_taxa(ps2, !Family == "Mitochondria")
sort(get_taxa_unique(ps3, taxonomic.rank = "Family"))
ps3
#Rarefy to 5000 reads
rngseed= 5
ps4 <- rarefy_even_depth(ps3, sample.size = 5000, rngseed = TRUE)

#Make relative abundance
ps4.rab <- transform_sample_counts(ps4, function(x) x/sum(x))

#Unifrac of the entire datasetps4.rab
psMTD <- ps4.rab
psMTD.df <- t(data.frame(otu_table(psMTD)))
psMTD.df <- data.frame(psMTD.df)
psMTD.df$abundance <- rowSums(psMTD.df[,c(1:180)])
psMTD.df$sequence <- rownames(psMTD.df)
psMTD.uniques <- getUniques(psMTD.df)
uniquesToFasta(psMTD.uniques, "MTD.fasta", ids = psMTD.df$sequence)
#After uniquesToFasta command, align seqs of the fasta file and then create phylogenetic tree. upload BestTree into read_tree command. For this publication I used CIPRES (available at https://www.phylo.org/portal2/login!input.action). Sequences were aligned using MAFFT on XSEDE (7.402), and phylogenetic tree made using RAxML-HPC BlackBox (8.2.12).
#Make Tree files
MTD.tree <- read_tree("~/MixedTrachyDataset/MTD_RAxML_bestTree.result")
psMTD.tree <- phyloseq(otu_table(otu_table(psMTD)),sample_data(sample_data(psMTD)), tax_table(tax_table(psMTD)), phy_tree(MTD.tree))

#Unweighted unifrac
MTD.uuf.dist <- distance(psMTD.tree, method = "unifrac")
MTD.uuf.ord <- ordinate(psMTD.tree, method = "PCoA", distance = "unifrac")
MTD.uuf.plot <- plot_ordination(psMTD.tree, MTD.uuf.ord, color = "State", shape = "Stage") +
scale_shape_manual(values = c(3,17,15,5,19))
plot(MTD.uuf.plot)

#PERMANOVA
psMTD.tree.data <- data.frame(sample_data(psMTD.tree))
adonis(MTD.uuf.dist ~State*Stage*Lab_or_Wild, psMTD.tree.data)

#Weighed Unifrac
MTD.wuf.dist <- distance(psMTD.tree, method = "wunifrac")
MTD.wuf.ord <- ordinate(psMTD.tree, method = "PCoA", distance = "wunifrac")
MTD.wuf.plot <- plot_ordination(psMTD.tree, MTD.wuf.ord, color = "State", shape = "Stage") + scale_shape_manual(values = c(3,17,15,5,19))
plot(MTD.wuf.plot)

#PERMANOVA
psMTD.tree.data <- data.frame(sample_data(psMTD.tree))
adonis(MTD.wuf.dist ~State*Stage*Lab_or_Wild, psMTD.tree.data)

#PERMANOVA
psMTD.tree.data <- data.frame(sample_data(psMTD.tree))
adonis(MTD.uuf.dist ~State*Stage*Colony.ID*Lab_or_Wild, psMTD.tree.data)

#Used the ASVs from the output to create phylogenetic tree.
write.csv(tax_table(ps4.rab), "MTD_ps4rab_taxtable.csv")
write.csv(otu_table(ps4.rab), "MTD_ps4rab_otutable.csv")

#Phylogenetic tree script
#Concatenated, ASVs of Mesoplasma and Spiroplasma, Mollicute strains from other fungus growing ants and Other strains the silva database to get the outline of the tree

cat MTD_ReferenceSeqs3.fasta TopMTD_Mollicute_Seqs.fasta SilvaMollicuteSpeciesRefined9.fasta > AllFiles10.fasta

#Align sequences
muscle -in AllFiles10.fasta -out AlignedAllFiles10.fasta
#Trimmed in MEGA-X (Molecular Evolutionary Genetics Analysis)
perl idk.phy AlignedAllFiles10Mega.fasta

#idk.phy file contains:
#!/usr/bin/perl -w
# obtained from Yu-Wei's Bioinformatics playground
# http://yuweibioinfo.blogspot.com/2009/01/fasta-to-phylip-converter.html
use strict;
MFAtoPHYLIP($ARGV[0]);
sub MFAtoPHYLIP
{
	my $inline;
	my $outfile = "$_[0]\.phy";
	my $count = 0;
	my $len;
	my $substate = 0;
	my @subheader;
	my @subcontent;
	my $m;
	my $n;
	open (FILE, "<$_[0]");
	while (defined($inline = <FILE>))
	{		chomp($inline);
		if ($inline =~ /^>([A-Za-z0-9.\-_:]+)/)
		{
			$subheader[$count] = $1;
			$subcontent[$count] = "";
			$count++;		}
		else
		{$subcontent[$count - 1] = $subcontent[$count - 1] . " $inline";
		}	}
	close (FILE);
	# Calculate the content length
	$n = length($subcontent[0]);
	$len = $n;
	for ($m = 0; $m < $n; $m++)
	{  if (substr($subcontent[0], $m, 1) eq " ")
		{   $len--;		}	}
	open (FILE, ">$outfile");
	print FILE "   $count    $len\n";
	for ($m = 0; $m < $count; $m++)
	{    $len = 10 - length($subheader[$m]);
		print FILE "$subheader[$m]";
		for ($n = 0; $n < $len; $n++)
		{	print FILE " ";	}
		print FILE " $subcontent[$m]\n";	}
	close (FILE);}

#Tree created with RAxMLHPC with 500 bootstraps
raxmlHPC -mGTRGAMMAI -s AlignedAllFiles10Mega.fasta.phy -n Feb20Tree -f a -N 500 -x 2 -p 5 -T 10
#Visualed the "besttree" in IToL (https://itol.embl.de/) and edited in adobe Illustrator

PUPAE DATASET
#Load dada2, I used v1.11.1  and followed their published workflow
library(dada2)
path <- "~/TrachyPupae/"
fnFs <- sort(list.files(path, pattern="_R1_001.fastq", full.name = TRUE))
fnRs <- sort(list.files(path, pattern="_R2_001.fastq", full.name = TRUE))
sample.names <- sapply(strsplit(basename(fnFs), "_"), `[`, 1)
#View quality plots and I ended up adjusted "truncLen" based on this.
plotQualityProfile(fnFs[1:2])
plotQualityProfile(fnRs[1:2])
filt_path <- file.path(path, "filtered")
filtFs <- file.path(filt_path, paste0(sample.names, "_F_filt.fastq"))
filtRs <- file.path(filt_path, paste0(sample.names, "_R_filt.fastq"))
out <- filterAndTrim(fnFs, filtFs, fnRs, filtRs, truncLen = c(240,160), maxN=0, maxEE = c(2,5), truncQ=2, rm.phix =TRUE, compress=TRUE, multithread = TRUE)
errF <- learnErrors(filtFs, multithread = TRUE)
errR <- learnErrors(filtRs, multithread = TRUE)
#View Errors
plotErrors(errF, nominalQ=TRUE)
derepFs <- derepFastq(filtFs, verbose=TRUE)
derepRs <- derepFastq(filtRs, verbose=TRUE)
names(derepFs) <- sample.names
names(derepRs) <- sample.names
dadaFs <- dada(derepFs, err=errF, multithread=TRUE)
dadaRs <- dada(derepRs, err=errR, multithread=TRUE)
dadaFs[[1]]
mergers <- mergePairs(dadaFs, derepFs, dadaRs, derepRs, verbose = TRUE)
seqtab <- makeSequenceTable(mergers)
seqtab.nochim <- removeBimeraDenovo(seqtab, method="consensus", multithread=TRUE, verbose = TRUE)
sum(seqtab.nochim)/sum(seqtab)
getN <- function(x) sum(getUniques(x))
track <- cbind(out, sapply(dadaFs, getN), sapply(mergers, getN), rowSums(seqtab), rowSums(seqtab.nochim))
colnames(track) <- c("input", "filtered", "denoised", "merged", "tabled", "nonchim")
rownames(track) <- sample.names
#Add in taxa information using your choice of database.
taxa <- assignTaxonomy(seqtab.nochim, "TrachyPupae/silva_nr_v128_train_set.fa", multithread = TRUE)
taxa.print <- taxa
rownames(taxa.print) <- NULL
head(taxa.print)
#Output the taxa, taxa counts, and taxa asv sequences
write.csv(taxa.print, "pupaetaxa.csv")
write.csv(seqtab.nochim, "pupaeASV.csv")

#Load other packages
library(phyloseq)
library(ggplot2)
library(plyr)
#Read in Metadata. Includes the sample name, whether it is a sample or control, and other information that is needed. I included whether they were ethanol washed.
meta <- read.csv("~/TrachyPupae/TrachyPupaeMetaData.csv",header = TRUE, row.names = 1)
ps <- phyloseq(otu_table(seqtab.nochim, taxa_are_rows = FALSE), sample_data(meta), tax_table(taxa))
#Check phyloseq object
ps
#Load decontam package
library(decontam)
sample_variables(ps)
df <- as.data.frame(sample_data(ps))
df$LibrarySize <- sample_sums(ps)
df <- df[order(df$LibrarySize),]
df$Index <- seq(nrow(df))
ggplot(data=df, aes(x=Index, y=LibrarySize, color=Sample_or_Control)) + geom_point()
sample_data(ps)$is.neg <- sample_data(ps)$Sample_or_Control == "Control Sample"
contamdf.prev <- isContaminant(ps, method="prevalence", neg="is.neg")
table(contamdf.prev$contaminant)

#To remove Eukyota, Archaea
ps1 <- subset_taxa(ps,Kingdom == "Bacteria")
#View it
get_taxa_unique(ps1, taxonomic.rank = "Kingdom")
#Remove anything not identified at Phylum level
ps2 <- subset_taxa(ps1, !is.na(Phylum))
sort(get_taxa_unique(ps2, taxonomic.rank = "Phylum"))
#Remove Mitochondria
ps3 <- subset_taxa(ps2, !Family == "Mitochondria")
sort(get_taxa_unique(ps3, taxonomic.rank = "Family"))
#ps4 rarefied to 10,000 reads
rngseed= 5
ps4 <- rarefy_even_depth(ps3, sample.size = 10000, rngseed = TRUE)

#Put into relative abundance
ps4.rab <- transform_sample_counts(ps4, function(x) x/sum(x))
#Glom asvs of the same genus together
ps4.rab.glom <- tax_glom(ps4.rab, taxrank = "Phylum")
ps4.rab.glom.df <- psmelt(ps4.rab.glom)
#Plot phylum level barplot
ps4.rab.glom.df$Phylum <- as.character(ps4.rab.glom.df$Phylum)
ps4.barplot.phylum <- ggplot(ps4.rab.glom.df, aes(x = Sample, y = Abundance, fill = Phylum, scale_color_brewer(pallette = "Set3"))) + geom_bar(stat = "identity") + theme(axis.text.x = element_text(angle = 270, vjust = 0.5))
plot(ps4.barplot.phylum)
#Arrange barplot by view by ethanol wash or not
ps4.barplot.phylum <- ggplot(ps4.rab.glom.df, aes(x = Sample, y = Abundance, fill = Phylum, scale_color_brewer(pallette = "Set3"))) + geom_bar(stat = "identity") + theme(axis.text.x = element_text(angle = 270, vjust = 0.5)) + facet_grid(~Wash, scales = "free", space = "free") + theme(strip.text = element_text(angle = 315), strip.background = element_blank(), panel.background = element_blank())
plot(ps4.barplot.phylum)
#Only include Phyla present at a 15% abundnace or higher, arranged by ethanol wash or not
max <- ddply(ps4.rab.glom.df, ~Phylum, function(x) c(max = max(x$Abundance)))
Other <- max[max$max <= 0.15,]$Phylum
ps4.rab.glom.df[ps4.rab.glom.df$Phylum %in% Other,]$Phylum <- "Other"
ps4.barplot.phylum <- ggplot(ps4.rab.glom.df, aes(x = Sample, y = Abundance, fill = Phylum, scale_color_brewer(pallette = "Set3"))) + geom_bar(stat = "identity") + theme(axis.text.x = element_text(angle = 270, vjust = 0.5)) + facet_grid(~Wash, scales = "free", space = "free") + theme(strip.text = element_text(angle = 315), strip.background = element_blank(), panel.background = element_blank())
plot(ps4.barplot.phylum)

#TopPhyla
ps4.barplot.phylum <- ggplot(ps4.rab.glom.df, aes(x = Sample, y = Abundance, fill = Phylum, scale_color_brewer(pallette = "Set3"))) + geom_bar(stat = "identity") + theme(axis.text.x = element_text(angle = 270, vjust = 0.5))
plot(ps4.barplot.phylum)

#psTener is looking at Tenericutes
psTener <- subset_taxa(ps4.rab, Phylum == "Tenericutes")
psTener.df <- psmelt(psTener)
psTener.df$Phylum <- as.character(psTener.df$Phylum)
psTener.barplot.phylum <- ggplot(psTener.df, aes(x = Sample, y = Abundance, fill = Genus, scale_color_brewer(pallette = "Set3"))) + geom_bar(stat = "identity") + theme(axis.text.x = element_text(angle = 270, vjust = 0.5))
plot(psTener.barplot.phylum)
psTener.barplot.phylum <- ggplot(psTener.df, aes(x = Sample, y = Abundance, fill = Genus, scale_color_brewer(pallette = "Set3"))) + geom_bar(stat = "identity") + theme(axis.text.x = element_text(angle = 270, vjust = 0.5)) + facet_grid(~Wash, scales = "free", space = "free") + theme(strip.text = element_text(angle = 315), strip.background = element_blank(), panel.background = element_blank())
#Look at relative abundance values and asv sequences
write.csv(psTener.df, "PupaeTenericutes.csv")


MULTISTATE DATASET
#Load dada2, I used v1.11.1
#I followed their published workflow and edited truncLen values.
library(dada2)
path <- "~/TrachyMicrobiome/"
fnFs <- sort(list.files(path, pattern="_R1_001.fastq", full.name = TRUE))
fnRs <- sort(list.files(path, pattern="_R2_001.fastq", full.name = TRUE))
sample.names <- sapply(strsplit(basename(fnFs), "_"), `[`, 1)
#Check quality
plotQualityProfile(fnFs[1:2])
plotQualityProfile(fnRs[1:2])
filt_path <- file.path(path, "filtered")
filtFs <- file.path(filt_path, paste0(sample.names, "_F_filt.fastq"))
filtRs <- file.path(filt_path, paste0(sample.names, "_R_filt.fastq"))
out <- filterAndTrim(fnFs, filtFs, fnRs, filtRs, truncLen = c(240,160), maxN=0, maxEE = c(2,5), truncQ=2, rm.phix =TRUE, compress=TRUE, multithread = TRUE)
errF <- learnErrors(filtFs, multithread = TRUE)
errR <- learnErrors(filtRs, multithread = TRUE)
#View errors
plotErrors(errF, nominalQ=TRUE)
derepFs <- derepFastq(filtFs, verbose=TRUE)
derepRs <- derepFastq(filtRs, verbose=TRUE)
names(derepFs) <- sample.names
names(derepRs) <- sample.names
dadaFs <- dada(derepFs, err=errF, multithread=TRUE)
dadaRs <- dada(derepRs, err=errR, multithread=TRUE)
dadaFs[[1]]
mergers <- mergePairs(dadaFs, derepFs, dadaRs, derepRs, verbose = TRUE)
seqtab <- makeSequenceTable(mergers)
seqtab.nochim <- removeBimeraDenovo(seqtab, method="consensus", multithread=TRUE, verbose = TRUE)
sum(seqtab.nochim)/sum(seqtab)
getN <- function(x) sum(getUniques(x))
track <- cbind(out, sapply(dadaFs, getN), sapply(mergers, getN), rowSums(seqtab), rowSums(seqtab.nochim))
colnames(track) <- c("input", "filtered", "denoised", "merged", "tabled", "nonchim")
rownames(track) <- sample.names
#Add in taxa information with database of choice
taxa <- assignTaxonomy(seqtab.nochim, "TrachyMicrobiome/silva_nr_v128_train_set.fa", multithread = TRUE)
taxa.print <- taxa
rownames(taxa.print) <- NULL
head(taxa.print)
#Output the taxa, taxa counts, and taxa asv sequences
write.csv(taxa.print, "taxa.csv")
write.csv(seqtab.nochim, "nameit.csv")

#Load other packages
library(phyloseq)
library(ggplot2)
library(plyr)
library(vegan)

#Read in Metadata. Includes the sample name, whether it is a sample or control, and other information that is needed. I included their state of location as well.
meta <- read.csv("~/TrachyMicrobiome/trachylearndeconmeta.csv",header = TRUE, row.names = 1)
ps <- phyloseq(otu_table(seqtab.nochim, taxa_are_rows = FALSE), sample_data(meta), tax_table(taxa))
#Check phyloseq object
ps
#Load decontam , to find if any ASVs are contamination.
library(decontam)
sample_variables(ps)
df <- as.data.frame(sample_data(ps))
df$LibrarySize <- sample_sums(ps)
df <- df[order(df$LibrarySize),]
df$Index <- seq(nrow(df))
ggplot(data=df, aes(x=Index, y=LibrarySize, color=Sample_or_Control)) + geom_point()
sample_data(ps)$is.neg <- sample_data(ps)$Sample_or_Control == "Control Sample"
contamdf.prev <- isContaminant(ps, method="prevalence", neg="is.neg")
table(contamdf.prev$contaminant)

#To remove Eukyota, Archaea
ps1 <- subset_taxa(ps,Kingdom == "Bacteria")
#View it
get_taxa_unique(ps1, taxonomic.rank = "Kingdom")
#remove Blank samples
ps2 <- subset_samples(ps1 , sample_names(ps1) != "Blank")
#Remove anything not identified at Phylum level
ps2 <- subset_taxa(ps2, !is.na(Phylum))
sort(get_taxa_unique(ps2, taxonomic.rank = "Phylum"))
#Remove Mitochondria
ps3 <- subset_taxa(ps2, !Family == "Mitochondria")
sort(get_taxa_unique(ps3, taxonomic.rank = "Family"))
#Double check phyloseq object
ps3
#Rarefy to even dataset
rngseed= 5
ps4 <- rarefy_even_depth(ps3, sample.size = 20000, rngseed = TRUE)

#Alpha Diversity violin plot. Using the same samples as ps4 after rarefying. "sample_sums" allows the same samples to be included without removing sequences.
ps3.5 <- subset_samples(ps3, sample_sums(ps3)>= 20000)
plot_richness(ps3.5, x = "State", measures = c("Shannon", "Simpson"), color = "Colony.ID")+ geom_violin() + theme(panel.background = element_blank(), panel.border = element_rect(fill = NA)) + stat_summary(fun.y= median, geom="point", shape=23, size=2)
plot_richness(ps3.5, x = "State", measures = c("Shannon", "Simpson"), color = "State")+ geom_violin() + theme(panel.background = element_blank(), panel.border = element_rect(fill = NA)) + stat_summary(fun.y= median, geom="point", shape=23, size=2)

#Create dataframe of alpha diversity values for anova stats
ps3.5richness.df <- estimate_richness(ps3.5, measures = c("Shannon", "Simpson"))
ps3.5richness.df$State <- sample_data(ps3.5)$State
ps3.5richness.df$Colony.ID <- sample_data(ps3.5)$Colony.ID
#Run anova on both Shannon and Simpson for state and colony id variables
ps3.5.state <- aov(Shannon ~State, ps3.5richness.df)
anova(ps3.5.state)
ps3.5.state <- aov(Simpson ~State, ps3.5richness.df)
anova(ps3.5.state)
ps3.5.colonyid <- aov(Shannon ~Colony.ID, ps3.5richness.df)
anova(ps3.5.colonyid)
ps3.5.colonyid <- aov(Simpson ~Colony.ID, ps3.5richness.df)
anova(ps3.5.colonyid)

#Phyla that are present
ps4.rab <- transform_sample_counts(ps4, function(x) x/sum(x))
ps4.rab.glom <- tax_glom(ps4.rab, taxrank = "Phylum")
ps4.rab.glom.df <- psmelt(ps4.rab.glom)
ps4.rab.glom.df$Phylum <- as.character(ps4.rab.glom.df$Phylum)
ps4.barplot.phylum <- ggplot(ps4.rab.glom.df, aes(x = Sample, y = Abundance, fill = Phylum, scale_color_brewer(pallette = "Set3"))) + geom_bar(stat = "identity") + theme(axis.text.x = element_text(angle = 270, vjust = 0.5)) + facet_grid(~State, scales = "free", space = "free") + theme(strip.text = element_text(angle = 315), strip.background = element_blank(), panel.background = element_blank())
plot(ps4.barplot.phylum)
#Top Phyla, present at 15% abundance or higher
max <- ddply(ps4.rab.glom.df, ~Phylum, function(x) c(max = max(x$Abundance)))
Other <- max[max$max <= 0.15,]$Phylum
ps4.rab.glom.df[ps4.rab.glom.df$Phylum %in% Other,]$Phylum <- "Other"

#Try To show top Genera
psTry <- ps4.rab
psTry.glom <- tax_glom(psTry, taxrank = "Genus")
psTry.glom.df <- psmelt(psTry.glom)
psTry.glom.df$Genus <- as.character(psTry.glom.df$Genus)
max <- ddply(psTry.glom.df, ~Genus, function(x) c(max = max(x$Abundance)))
Other <- max[max$max <= 0.15,]$Genus
psTry.glom.df[psTry.glom.df$Genus %in% Other,]$Genus <- "Other"
psTry.barplot.genus <- ggplot(psTry.glom.df, aes(x = Sample, y = Abundance, fill = Genus, scale_color_brewer(pallette = "Set3"))) + geom_bar(stat = "identity") + theme(axis.text.x = element_text(angle = 270, vjust = 0.5)) + facet_grid(~State, scales = "free", space = "free") + theme(strip.text = element_text(angle = 315), strip.background = element_blank(), panel.background = element_blank())
plot(psTry.barplot.genus)

#ps5- Tenericutes
ps5 <- subset_taxa(ps4.rab, Phylum == "Tenericutes")
ps5.df <- psmelt(ps5)
ps5.df$Phylum <- as.character(ps5.df$Phylum)
ps5.barplot.phylum <- ggplot(ps5.df, aes(x = Sample, y = Abundance, fill = Genus, scale_color_brewer(pallette = "Set3"))) + geom_bar(stat = "identity") + theme(axis.text.x = element_text(angle = 270, vjust = 0.5))
plot(ps5.barplot.phylum)
ps5.barplot.phylum <- ggplot(ps5.df, aes(x = Sample, y = Abundance, fill = Phylum, scale_color_brewer(pallette = "Set3"))) + geom_bar(stat = "identity") + theme(axis.text.x = element_text(angle = 270, vjust = 0.5)) + facet_grid(~State, scales = "free", space = "free") + theme(strip.text = element_text(angle = 315), strip.background = element_blank(), panel.background = element_blank())

#BetaDiversity Tests
#All States
ps4.df <- t(data.frame(otu_table(ps4)))
ps4.df <- data.frame(ps4.df)
ps4.df$abundance <- rowSums(ps4.df[,(1:71)])
ps4.df$sequence <- rownames(ps4.df)
ps4.uniques <- getUniques(ps4.df)
uniquesToFasta(ps4.uniques, "AllStates.fasta", ids = ps4.df$sequence)
#After uniquesToFasta command, align seqs of the fasta file and then create phylogenetic tree. upload BestTree into read_tree command. For this publication I used CIPRES (available at https://www.phylo.org/portal2/login!input.action). Sequences were aligned using MAFFT on XSEDE (7.402), and phylogenetic tree made using RAxML-HPC BlackBox (8.2.12).

#Make Tree files
AllStates.tree <- read_tree("~/TrachyMicrobiome/AllStatesRAxML_bestTree.result")
ps4.tree <- phyloseq(otu_table(otu_table(ps4)), sample_data(sample_data(ps4)), tax_table(tax_table(ps4)), phy_tree(AllStates.tree))
#Unweighted unifrac
ps4.uuf.dist <- distance(ps4.tree, method = "unifrac")
ps4.uuf.ord <- ordinate(ps4.tree, method = "PCoA", distance = "unifrac")
ps4.uuf.plot <- plot_ordination(ps4.tree, ps4.uuf.ord, color = "State")
plot(ps4.uuf.plot) + stat_ellipse()
#PERMANOVA
ps4.tree.data <- data.frame(sample_data(ps4.tree))
adonis(ps4.uuf.dist ~State*Colony.ID, ps4.tree.data)

#Weighted unifrac
ps4.wuf.dist <- distance(ps4.tree, method = "wunifrac")
ps4.wuf.ord <- ordinate(ps4.tree, method = "PCoA", distance = "wunifrac")
ps4.wuf.plot <- plot_ordination(ps4.tree, ps4.wuf.ord, color = "State")
plot(ps4.wuf.plot) + stat_ellipse()
#PERMANOVA
ps4.tree.data <- data.frame(sample_data(ps4.tree))
adonis(ps4.wuf.dist ~ State*Colony.ID, ps4.tree.data)

#Remove NC & NY
psRemove <- subset_samples(ps4.rab, !sample_data(ps4.rab)$State == "North Carolina")
psRemove <- subset_samples(psRemove, !sample_data(psRemove)$State == "New York")
psRemove.df <- t(data.frame(otu_table(psRemove)))
psRemove.df <- data.frame(psRemove.df)
psRemove.df$abundance <- rowSums(psRemove.df[,(1:52)])
psRemove.df$sequence <- rownames(psRemove.df)
psRemove.uniques <- getUniques(psRemove.df)
uniquesToFasta(psRemove.uniques, "NoNCNY.fasta", ids = psRemove.df$sequence)
#After uniquesToFasta command, align seqs of the fasta file and then create phylogenetic tree. upload BestTree into read_tree command. For this publication I used CIPRES (available at https://www.phylo.org/portal2/login!input.action). Sequences were aligned using MAFFT on XSEDE (7.402), and phylogenetic tree made using RAxML-HPC BlackBox (8.2.12).
#Make Tree files
Remove.tree <- read_tree("~/TrachyMicrobiome/NoNYorNC_bestTree.result")
psRemove.tree <- phyloseq(otu_table(otu_table(psRemove)), sample_data(sample_data(psRemove)), tax_table(tax_table(psRemove)), phy_tree(Remove.tree))

#Unweighted unifrac
psRemove.uuf.dist <- distance(psRemove.tree, method = "unifrac")
psRemove.uuf.ord <- ordinate(psRemove.tree, method = "PCoA", distance = "unifrac")
psRemove.uuf.plot <- plot_ordination(psRemove.tree, psRemove.uuf.ord, color = "State")
plot(psRemove.uuf.plot)
#PERMANOVA
psRemove.tree.data <- data.frame(sample_data(psRemove.tree))
adonis(psRemove.uuf.dist ~State*Colony.ID, psRemove.tree.data)

#Weighted unifrac
psRemove.wuf.dist <- distance(psRemove.tree, method = "wunifrac")
psRemove.wuf.ord <- ordinate(psRemove.tree, method = "PCoA", distance = "wunifrac")
psRemove.wuf.plot <- plot_ordination(psRemove.tree, psRemove.wuf.ord, color = "State")
plot(psRemove.wuf.plot)
#PERMANOVA
psRemove.tree.data <- data.frame(sample_data(psRemove.tree))
adonis(psRemove.wuf.dist ~State*Colony.ID, psRemove.tree.data)

#MANTEL TEST & Correlog!
#Only include one sample from each colony for this section.
#Create a csv with 3 rows titled: sample name, latitude and longitude. Fill in with corresponding data. Upload csv into GeoMatrix (Ersts, 2021).
#*Ersts, P.J [Internet] Geographic Distance Matrix Generator (version 1.2.3). American Museum of Natural History, Center for Biodiversity and Conservation. Available from http://biodiversityinformatics.amnh.org/open_source/gdmg.  Accessed on 2021-8-27.

geodist.Edit.matrix <- read.delim("~/TrachyMicrobiome/GeoMatrixOutputEdit.txt", header = TRUE, row.names = 1, check.names = FALSE)
geodist.Edit.matrix <- as.matrix(geodist.Edit.matrix)

#Use weighted unifrac distance matrix from above
ps4.wuf.dist.matrix <- as.matrix(ps4.wuf.dist)
write.csv(ps4.wuf.dist.matrix, "ps4.wuf.dist.csv")
#Removed samples from the same colony and saved file as a table delimited text file. Uploaded back into R
ps4.edit.wuf.dist.matrix <- read.table("~/TrachyMicrobiome/ps4.Edit.wuf.dist.txt")
ps4.edit.wuf.dist.matrix <- as.matrix(ps4.edit.wuf.dist.matrix)

#Mantel test
mantel(ps4.edit.wuf.dist.matrix,geodist.Edit.matrix)
#Plot Mantel correlogram
ps4.Edit.wuf.corelog <- mantel.correlog(ps4.edit.wuf.dist.matrix,geodist.Edit.matrix)
plot(ps4.Edit.wuf.corelog)
#Partial mantel stats
mantel.correlog(ps4.edit.wuf.dist.matrix,geodist.Edit.matrix)

#Use unweighted unifrac distance matrix from above
ps4.uuf.dist.matrix <- as.matrix(ps4.uuf.dist)
write.csv(ps4.uuf.dist.matrix, "ps4.uuf.dist.csv")
#Removed samples from the same colony and saved file as a table delimited text file. Uploaded back into R
ps4.edit.uuf.dist.matrix <- read.table("~/TrachyMicrobiome/ps4.Edit.uuf.dist.txt")
ps4.edit.uuf.dist.matrix <- as.matrix(ps4.edit.uuf.dist.matrix)

#Mantel test
mantel(ps4.edit.uuf.dist.matrix,geodist.Edit.matrix)
#Plot Mantel correlogram
ps4.Edit.uuf.corelog <- mantel.correlog(ps4.edit.uuf.dist.matrix, geodist.Edit.matrix)
plot(ps4.Edit.uuf.corelog)
#Partial mantel stats
mantel.correlog(ps4.edit.uuf.dist.matrix, geodist.Edit.matrix)
#Only include genus if it is present at least once time at 5% or greater. All others get pushed into "Other"
psTry <- ps4.rab
psTry.glom <- tax_glom(psTry, taxrank = "Genus")
psTry.glom.df <- psmelt(psTry.glom)
psTry.glom.df$Genus <- as.character(psTry.glom.df$Genus)
max <- ddply(psTry.glom.df, ~Genus, function(x) c(max = max(x$Abundance)))
Other <- max[max$max <= 0.05,]$Genus
psTry.glom.df[psTry.glom.df$Genus %in% Other,]$Genus <- "Other"
psTry.barplot.genus <- ggplot(psTry.glom.df, aes(x = Sample, y = Abundance, fill = Genus, scale_color_brewer(pallette = "Set3"))) + geom_bar(stat = "identity") + theme(axis.text.x = element_text(angle = 270, vjust = 0.5)) + facet_grid(~State, scales = "free", space = "free") + theme(strip.text = element_text(angle = 315), strip.background = element_blank(), panel.background = element_blank())
plot(psTry.barplot.genus)

#Need OTU table and taxa output to merge counts. ASVs  on the left side, Samples across the top
write.csv(tax_table(psTry.glom), "MultiStateTaxTableHeatmapPsTry5.csv")
#ABUNDANCE DOT PLOT & BOX PLOT  IT WORKED!
psTry.dotplot.genus <- ggplot(psTry.glom.df, aes(x = Genus, y = Abundance)) + geom_point(aes(fill= State, color = State), alpha = 0.5) + theme(axis.text.x = element_text(angle = 270, vjust = 0.5)) + theme(strip.text = element_text(angle = 315), strip.background = element_blank(), panel.background = element_blank())
plot(psTry.dotplot.genus)
psTry.boxplot.genus <- ggplot(psTry.glom.df, aes(x = Genus, y = Abundance)) + geom_boxplot(outlier.alpha = 0.2) + theme(axis.text.x = element_text(angle = 270, vjust = 0.5)) + theme(strip.text = element_text(angle = 315), strip.background = element_blank(), panel.background = element_blank())
plot(psTry.boxplot.genus)

#Took the "MultiStateTaxTableHeatmapPsTry5.csv" and edited in excel. Need taxa as rows. In excel, calculate prevalence and median relatative abudnance of the taxa
#Import back into R
prevalencedf <- read.csv("~/TrachyMicrobiome/PrevalenceMultiStateHeatmap18taxaPsTry5.csv")
#Bubble chart, Prevalence in x axis, median relative abundance is the bubble shape and genus name is on the y axis.  Prevalence by genus & bubble is the RAB
prevalencedf.plot <- ggplot(prevalencedf, aes(x= reorder(Genus, -PercentPrevalence.5.AbundDataset), y= PercentPrevalence.5.AbundDataset, size = MedianRelAbund)) + geom_point(aes(fill= MedianRelAbund)) + theme(axis.text.x = element_text(angle = 270, vjust = 0.5)) + theme(strip.text = element_text(angle = 315), strip.background = element_blank(), panel.background = element_blank())
plot(prevalencedf.plot)

Two-Colony Dataset
#Figure 4C was created by exporting the WUF and UUF distance files into excel and calculating the median. A heatmap was created in excel and edited in adobe illustrator.
library(dada2)
path <- "~/AntDec2018/"
fnFs <- sort(list.files(path, pattern="_R1_001.fastq", full.name = TRUE))
fnRs <- sort(list.files(path, pattern="_R2_001.fastq", full.name = TRUE))
sample.names <- sapply(strsplit(basename(fnFs), "_"), `[`, 1)
plotQualityProfile(fnFs[1:2])
plotQualityProfile(fnRs[1:2])
#View the read quality
filt_path <- file.path(path, "filtered")
filtFs <- file.path(filt_path, paste0(sample.names, "_F_filt.fastq"))
filtRs <- file.path(filt_path, paste0(sample.names, "_R_filt.fastq"))
out <- filterAndTrim(fnFs, filtFs, fnRs, filtRs, truncLen = c(240,160), maxN=0, maxEE = c(2,5), truncQ=2, rm.phix =TRUE, compress=TRUE, multithread = TRUE)
errF <- learnErrors(filtFs, multithread = TRUE)
errR <- learnErrors(filtRs, multithread = TRUE)
plotErrors(errF, nominalQ=TRUE)
#Look at errors
derepFs <- derepFastq(filtFs, verbose=TRUE)
derepRs <- derepFastq(filtRs, verbose=TRUE)
names(derepFs) <- sample.names
names(derepRs) <- sample.names
dadaFs <- dada(derepFs, err=errF, multithread=TRUE)
dadaRs <- dada(derepRs, err=errR, multithread=TRUE)
dadaFs[[1]]
mergers <- mergePairs(dadaFs, derepFs, dadaRs, derepRs, verbose = TRUE)
seqtab <- makeSequenceTable(mergers)
seqtab.nochim <- removeBimeraDenovo(seqtab, method="consensus", multithread=TRUE, verbose = TRUE)
sum(seqtab.nochim)/sum(seqtab)
getN <- function(x) sum(getUniques(x))
track <- cbind(out, sapply(dadaFs, getN), sapply(mergers, getN), rowSums(seqtab), rowSums(seqtab.nochim))
colnames(track) <- c("input", "filtered", "denoised", "merged", "tabled", "nonchim")
rownames(track) <- sample.names
#Add in database of choice for taxonomy
taxa <- assignTaxonomy(seqtab.nochim, "AntDec2018/silva_nr_v128_train_set.fa", multithread = TRUE)
taxa.print <- taxa
rownames(taxa.print) <- NULL
head(taxa.print)
write.csv(taxa.print, "TaxaListAntDec2018.csv")
write.csv(seqtab.nochim, "NameListAntDec2018.csv")

library(phyloseq)
library(ggplot2)
library(plyr)
library(vegan)
#Read in Metadata, I included the caste type, sample setting and colony ID.
meta <- read.csv("~/AntDec2018/MetaDataAntDec18.csv",header = TRUE, row.names = 1)
ps <- phyloseq(otu_table(seqtab.nochim, taxa_are_rows = FALSE), sample_data(meta), tax_table(taxa))
ps
#Decontam will help determine if there are ASVs shared between the negative controls and samples and lets you remove the ASVs flagged as contamination
library(decontam)
sample_variables(ps)
df <- as.data.frame(sample_data(ps))
df$LibrarySize <- sample_sums(ps)
df <- df[order(df$LibrarySize),]
df$Index <- seq(nrow(df))
ggplot(data=df, aes(x=Index, y=LibrarySize, color=Sample_or_Control)) + geom_point()
sample_data(ps)$is.neg <- sample_data(ps)$Sample_or_Control == "Control Sample"
contamdf.prev <- isContaminant(ps, method="prevalence", neg="is.neg")
table(contamdf.prev$contaminant)
##To remove Eukyota, Archaea
ps1 <- subset_taxa(ps,Kingdom == "Bacteria")
#View it
get_taxa_unique(ps1, taxonomic.rank = "Kingdom")
#Remove anything not identified at Phylum level
ps2 <- subset_taxa(ps2, !is.na(Phylum))
sort(get_taxa_unique(ps2, taxonomic.rank = "Phylum"))
#Remove Mitochondria
ps3 <- subset_taxa(ps2, !Family == "Mitochondria")
sort(get_taxa_unique(ps3, taxonomic.rank = "Family"))
ps3
#Remove Blanks & Water, need to use the sample name.
ps3 <- subset_samples(ps3, sample_names(ps3) != "NFH2O-712-508")
#ps3 has 147 samples
#This allows for Alpha diversity on non transformed/rarefied data.
#Includes only samples that have 5000 reads or more.
ps3.5 <- subset_samples(ps3, sample_sums(ps3)>= 5000)
psRAW270 <- subset_samples(ps3.5, sample_data(ps3.5)$Colony.ID == "JKH000270")
psRAW307 <- subset_samples(ps3.5, sample_data(ps3.5)$Colony.ID == "JKH000307")

#Alpha Diversity
#JKH000270
plot_richness(psRAW270, x = "Stage", measures = c("Shannon", "Simpson"), color = "Lab_or_Wild")
plot_richness(psRAW270, x = "Lab_or_Wild", measures = c("Shannon", "Simpson"), color = "Stage")
#JKH000307
plot_richness(psRAW307, x = "Stage", measures = c("Shannon", "Simpson"), color = "Lab_or_Wild")
plot_richness(psRAW307, x = "Lab_or_Wild", measures = c("Shannon", "Simpson"), color = "Stage")

#Alpha div. violin plot
plot_richness(psRAW270, x = "Stage", measures = c("Shannon", "Simpson"), color = "Lab_or_Wild")+ geom_violin() + theme(panel.background = element_blank(), panel.border = element_rect(fill = NA)) + stat_summary(fun.y= median, geom="point", shape=23, size=2)
plot_richness(psRAW307, x = "Stage", measures = c("Shannon", "Simpson"), color = "Lab_or_Wild")+ geom_violin() + theme(panel.background = element_blank(), panel.border = element_rect(fill = NA)) + stat_summary(fun.y= median, geom="point", shape=23, size=2)

#Stats for comparing the alpha diversitys between the colony, caste and lab adaptation
library(dunn.test)
#Entire dataset, compare by Colony ID
TwoColAlphaDiv.df <- estimate_richness(ps3.5, measures = c("Simpson","Shannon"))
TwoColAlphaDiv.df$Colony.ID <-sample_data(ps3.5)$Colony.ID
dunn.test(TwoColAlphaDiv.df$Shannon, g = TwoColAlphaDiv.df$Colony.ID, method = "bonferroni")
dunn.test(TwoColAlphaDiv.df$Simpson, g = TwoColAlphaDiv.df$Colony.ID, method = "bonferroni")

#Multiple test corrections on the alpha diversity data
#Using anova test
ps3.5richness.df <- estimate_richness(ps3.5, measures = c("Shannon", "Simpson"))
ps3.5richness.df$Stage <- sample_data(ps3.5)$Stage
ps3.5richness.df$Colony.ID <- sample_data(ps3.5)$Colony.ID
ps3.5richness.df$Lab_or_Wild <- sample_data(ps3.5)$Lab_or_Wild
ps3.5.colonyid <- aov(Shannon ~Colony.ID, ps3.5richness.df)
anova(ps3.5.colonyid)

ps3.5.colonyid <- aov(Simpson ~Colony.ID, ps3.5richness.df)
anova(ps3.5.colonyid)
ps3.5.stage <- aov(Shannon ~Stage, ps3.5richness.df)
anova(ps3.5.stage)
TukeyHSD(ps3.5.stage)
ps3.5.stage <- aov(Simpson ~Stage, ps3.5richness.df)
anova(ps3.5.stage)
ps3.5.low <- aov(Shannon ~Lab_or_Wild, ps3.5richness.df)
anova(ps3.5.low)
ps3.5.low <- aov(Simpson ~Lab_or_Wild, ps3.5richness.df)
anova(ps3.5.low)
ps3.5.all <- aov(Shannon ~Colony.ID*Stage*Lab_or_Wild, ps3.5richness.df)
anova(ps3.5.all)
ps3.5.all <- aov(Simpson ~Colony.ID*Stage*Lab_or_Wild, ps3.5richness.df)
anova(ps3.5.all)

#Rarefy to even number.
rngseed= 5
ps4 <- rarefy_even_depth(ps3, sample.size = 5000, rngseed = TRUE)

#Phyla that are present in entire dataset
ps4.rab <- transform_sample_counts(ps4, function(x) x/sum(x))
ps4.rab.glom <- tax_glom(ps4.rab, taxrank = "Phylum")
ps4.rab.glom.df <- psmelt(ps4.rab.glom)
ps4.rab.glom.df$Phylum <- as.character(ps4.rab.glom.df$Phylum)
ps4.barplot.phylum <- ggplot(ps4.rab.glom.df, aes(x = Sample, y = Abundance, fill = Phylum, scale_color_brewer(pallette = "Set3"))) + geom_bar(stat = "identity") + theme(axis.text.x = element_text(angle = 270, vjust = 0.5)) + facet_grid(~Colony.ID, scales = "free", space = "free") + theme(strip.text = element_text(angle = 315), strip.background = element_blank(), panel.background = element_blank())
plot(ps4.barplot.phylum)
#Top Phyla
max <- ddply(ps4.rab.glom.df, ~Phylum, function(x) c(max = max(x$Abundance)))
Other <- max[max$max <= 0.15,]$Phylum
ps4.rab.glom.df[ps4.rab.glom.df$Phylum %in% Other,]$Phylum <- "Other"

#Genera
ps5 <- tax_glom(ps4.rab, taxrank = "Genus")
ps5.glom.df <- psmelt(ps5)
ps5.glom.df$Genus <- as.character(ps5.glom.df$Genus)
max <- ddply(ps5.glom.df, ~Genus, function(x) c(max = max(x$Abundance)))
Other <- max[max$max <= 0.15,]$Genus
ps5.glom.df[ps5.glom.df$Genus %in% Other,]$Genus <- "Other"
ps5.barplot.genus <- ggplot(ps5.glom.df, aes(x = Sample, y = Abundance, fill = Genus, scale_color_brewer(pallette = "Set3"))) + geom_bar(stat = "identity") + theme(axis.text.x = element_text(angle = 270, vjust = 0.5)) + facet_grid(~Colony.ID, scales = "free", space = "free") + theme(strip.text = element_text(angle = 315), strip.background = element_blank(), panel.background = element_blank())
plot(ps5.barplot.genus)
#Phyloseq for just JKH270
ps270 <- subset_samples(ps4.rab, sample_data(ps4.rab)$Colony.ID == "JKH000270")
#All Phyla
ps270.glom <- tax_glom(ps270, taxrank = "Phylum")
ps270.glom.df <- psmelt(ps270.glom)
ps270.glom.df$Phylum <- as.character(ps270.glom.df$Phylum)
ps270.barplot.phylum <- ggplot(ps270.glom.df, aes(x = Sample, y = Abundance, fill = Phylum, scale_color_brewer(pallette = "Set3"))) + geom_bar(stat = "identity") + theme(axis.text.x = element_text(angle = 270, vjust = 0.5)) + facet_grid(~Stage, scales = "free", space = "free") + theme(strip.text = element_text(angle = 315), strip.background = element_blank(), panel.background = element_blank())
plot(ps270.barplot.phylum)
#Top Phyla
max <- ddply(ps270.glom.df, ~Phylum, function(x) c(max = max(x$Abundance)))
Other <- max[max$max <= 0.15,]$Phylum
ps270.glom.df[ps270.glom.df$Phylum %in% Other,]$Phylum <- "Other"

#ps270Ten Tenericutes
ps270Ten <- subset_taxa(ps270, Phylum == "Tenericutes")
ps270Ten.df <- psmelt(ps270Ten)
ps270Ten.df$Phylum <- as.character(ps270Ten.df$Phylum)

ps270Ten.barplot.phylum <- ggplot(ps270Ten.df, aes(x = Sample, y = Abundance, fill = Genus, scale_color_brewer(pallette = "Set3"))) + geom_bar(stat = "identity") + theme(axis.text.x = element_text(angle = 270, vjust = 0.5)) + facet_grid(~Stage, scales = "free", space = "free") + theme(strip.text = element_text(angle = 315), strip.background = element_blank(), panel.background = element_blank())
plot(ps270Ten.barplot.phylum)
write.csv(ps270Ten.df, "270Tenericutes.csv")
write.csv(ps4.rab, "AntDecRelativeAbundance")
write.csv(ps4.rab.glom.df, "Trial.csv")

#Phyloseq for JKH307
ps307 <- subset_species(ps4.rab, sample_data(ps4.rab)$Colony.ID == "JKH000307")
#All Phyla
ps307.glom <- tax_glom(ps307, taxrank = "Phylum")
ps307.glom.df <- psmelt(ps307.glom)
ps307.glom.df$Phylum <- as.character(ps307.glom.df$Phylum)
ps307.barplot.phylum <- ggplot(ps307.glom.df, aes(x = Sample, y = Abundance, fill = Phylum, scale_color_brewer(pallette = "Set3"))) + geom_bar(stat = "identity") + theme(axis.text.x = element_text(angle = 270, vjust = 0.5)) + facet_grid(~Stage, scales = "free", space = "free") + theme(strip.text = element_text(angle = 315), strip.background = element_blank(), panel.background = element_blank())
plot(ps307.barplot.phylum)

#Top Phyla
max <- ddply(ps307.glom.df, ~Phylum, function(x) c(max = max(x$Abundance)))
Other <- max[max$max <= 0.15,]$Phylum
ps307.glom.df[ps307.glom.df$Phylum %in% Other,]$Phylum <- "Other"
#ps307Ten Tenericutes
ps307Ten <- subset_taxa(ps307, Phylum == "Tenericutes")
ps307Ten.df <- psmelt(ps307Ten)
ps307Ten.df$Phylum <- as.character(ps307Ten.df$Phylum)
ps307Ten.barplot.phylum <- ggplot(ps307Ten.df, aes(x = Sample, y = Abundance, fill = Genus, scale_color_brewer(pallette = "Set3"))) + geom_bar(stat = "identity") + theme(axis.text.x = element_text(angle = 270, vjust = 0.5)) + facet_grid(~Stage, scales = "free", space = "free") + theme(strip.text = element_text(angle = 315), strip.background = element_blank(), panel.background = element_blank())
plot(ps307Ten.barplot.phylum)
write.csv(ps307Ten.df, "307Tenericutes.csv")

#HeatMap
library(gplots)
#Create a top genus barplot to find the top genera,then create heatmap
psTry <- ps4.rab
psTry.glom <- tax_glom(psTry, taxrank = "Genus")
psTry.glom.df <- psmelt(psTry.glom)
psTry.glom.df$Genus <- as.character(psTry.glom.df$Genus)
max <- ddply(psTry.glom.df, ~Genus, function(x) c(max = max(x$Abundance)))
Other <- max[max$max <= 0.05,]$Genus
psTry.glom.df[psTry.glom.df$Genus %in% Other,]$Genus <- "Other"
psTry.barplot.genus <- ggplot(psTry.glom.df, aes(x = Sample, y = Abundance, fill = Genus, scale_color_brewer(pallette = "Set3"))) + geom_bar(stat = "identity") + theme(axis.text.x = element_text(angle = 270, vjust = 0.5)) + facet_grid(~Colony.ID*Stage*Lab_or_Wild, scales = "free", space = "free") + theme(strip.text = element_text(angle = 315), strip.background = element_blank(), panel.background = element_blank())
plot(psTry.barplot.genus)

#Need OTU table and taxa output to merge counts. ASVs  on the left side, Samples across the top
#taxa table
write.csv(tax_table(psTry.glom), "TwoColonyTaxTablePsTry5.csv")
#otu counts table
write.csv(otu_table(psTry.glom), "TwoColonyOTUTablePsTry5.csv")
#I took the top genera that were from the barplot... I couldn't figure out how to do it any other way.
#Run entire script to the barplot to find what taxa need to be included.
#A genus is only present if it was found to be at a 5% abudance or greater, in any sample.
hmdata3 <- read.csv("~/AntDec2018/AntDec2018kek/TwoColonyHeatmapPsTry5.csv", header=TRUE, row.names = 1, sep = ",")
hmdata3 <- as.matrix(hmdata3[,-dim(hmdata3)[2]])
madTry <- apply(hmdata3, 1, sum)
madTry <- sort(madTry, decreasing = TRUE)
madTry <- madTry[1:26]
madTrytaxa <- hmdata3[row.names(hmdata3) %in% names(madTry),]
Bluecolor = colorRampPalette(colors = c("white","royalblue3"))
heatmap.2(as.matrix(madTrytaxa), trace = "none", col = Bluecolor, cexCol = 0.3, cexRow = 0.75, distfun = function(x) dist(x, method = "euclidean"), hclustfun = function(x) hclust(x, method="ward.D"), margins = c(6,6), key = TRUE, keysize = 2.0)

#Script to edit shapes
scale_shape_manual(values = c(3,17,15,16))
#The caste was always shown in alphabetical order, so this helps keep shapes the same with the order of caste.

#Both colonies
psAll <- ps4.rab
psAll.df <- t(data.frame(otu_table(psAll)))
psAll.df <- data.frame(psAll.df)
psAll.df$abundance <- rowSums(psAll.df[,c(1:110)])
psAll.df$sequence <- rownames(psAll.df)
psAll.uniques <- getUniques(psAll.df)
uniquesToFasta(psAll.uniques, "BothJKHs.fasta", ids = psAll.df$sequence)
#After uniquesToFasta command, align seqs of the fasta file and then create phylogenetic tree. upload BestTree into read_tree command. For this publication I used CIPRES (available at https://www.phylo.org/portal2/login!input.action). Sequences were aligned using MAFFT on XSEDE (7.402), and phylogenetic tree made using RAxML-HPC BlackBox (8.2.12).
#Make Tree files
All.tree <- read_tree("~/AntDec2018/AntDec2018kek/BothColonyRAxML_bestTree.result")
psAll.tree <- phyloseq(otu_table(otu_table(psAll)),sample_data(sample_data(psAll)), tax_table(tax_table(psAll)), phy_tree(All.tree))

#Unweighted unifrac
All.uuf.dist <- distance(psAll.tree, method = "unifrac")
All.uuf.ord <- ordinate(psAll.tree, method = "PCoA", distance = "unifrac")
All.uuf.plot <- plot_ordination(psAll.tree, All.uuf.ord, color = "Stage", shape = "Lab_or_Wild")
plot(All.uuf.plot)
plot(All.uuf.plot) + stat_ellipse(aes(group = Colony.ID))
All.uuf.plot <- plot_ordination(psAll.tree, All.uuf.ord, color = "Colony.ID", shape = "Lab_or_Wild")
plot(All.uuf.plot)
All.uuf.plot <- plot_ordination(psAll.tree, All.uuf.ord, color = "Stage", shape = "Colony.ID")
plot(All.uuf.plot)
#PERMANOVA
psAll.tree.data <- data.frame(sample_data(psAll.tree))
adonis(All.uuf.dist ~Stage*Lab_or_Wild*Colony.ID, psAll.tree.data)

#Weighed Unifrac
All.wuf.dist <- distance(psAll.tree, method = "wunifrac")
All.wuf.ord <- ordinate(psAll.tree, method = "PCoA", distance = "wunifrac")
All.wuf.plot <- plot_ordination(psAll.tree, All.wuf.ord, color = "Stage", shape = "Lab_or_Wild")
plot(All.wuf.plot)
plot(All.wuf.plot) + stat_ellipse(aes(group = Colony.ID))
All.wuf.plot <- plot_ordination(psAll.tree, All.wuf.ord, color = "Colony.ID", shape = "Lab_or_Wild")
plot(All.wuf.plot)
All.wuf.plot <- plot_ordination(psAll.tree, All.wuf.ord, color = "Stage", shape = "Colony.ID")
plot(All.wuf.plot)
#PERMANOVA
psAll.tree.data <- data.frame(sample_data(psAll.tree))
adonis(All.wuf.dist ~Stage*Lab_or_Wild*Colony.ID, psAll.tree.data)


#Stats for Suppl. Fig 11
#Make vectors of all the weighted and unweighted ant data so that
#I can do a KS test to check if Data is normalized. If normalized then I can do
# a t-test in R, or non normalized I will use a MannWhitney test
Malates307UWild <- c(0.820192504, 0.816284355, 0.904745997, 0.818555097, 0.843629643, 0.800818592, 0.372750955, 0.354876672, 0.601040305, 0.343448414, 0.43789047, 0.488119031, 0.47087094, 0.464447201, 0.41594612, 0.418163795, 0.420549305, 0.578134287, 0.596019971, 0.571163302, 0.497216184)
Malates307ULab <- c(0.265793972, 0.258864185, 0.27867304, 0.276625371, 0.355723507, 0.261717003, 0.30467295, 0.347363613, 0.299232086, 0.319677318, 0.33521439, 0.309638392, 0.302049376, 0.304686974, 0.27441542, 0.064595626, 0.211021379, 0.131181081, 0.128892715, 0.110711976, 0.108834123, 0.347564689, 0.092317394, 0.125891502, 0.154262453, 0.110767489, 0.160004957, 0.15490151, 0.207785596, 0.202510451, 0.121628201, 0.163986482, 0.100701074, 0.104953206, 0.35742127, 0.106622318, 0.131696211, 0.146365809, 0.124608962, 0.15075875, 0.168404445, 0.199420115, 0.154640869, 0.233191943, 0.133859954, 0.16956236, 0.372298811, 0.185846908, 0.150749663, 0.208790101, 0.197420118, 0.126561234, 0.123447346, 0.182382863, 0.211830717, 0.025348438, 0.116824154, 0.333166061, 0.041193115, 0.096724923, 0.162823776, 0.080402711, 0.033965159, 0.127059208, 0.123878707, 0.194413124, 0.219414902, 0.363218638, 0.175410392, 0.203380731, 0.234569026, 0.137487025, 0.188315697, 0.226847589, 0.280982179, 0.137249068,	0.348758253,	0.063281366,	0.117593196,	0.182230407,	0.101630324,	0.056966936,	0.147915842,	0.1455977,	0.289697512,	0.078964695,	0.046161811,	0.079068238,	0.097337949,	0.144876495, 0.075601003,	0.219074671,	0.304762686,	0.277863441,	0.317322885,	0.278041896,	0.354515766,	0.299799181,	0.402673508,	0.057980944,	0.126984813,	0.040939677,	0.071562729,	0.088431384,	0.156001323,	0.109572651,	0.102614697,	0.188629831,	0.181885416,	0.15570032,	0.153705391,	0.139185248,	0.189460162, 0.097665019,	0.257715206,	0.095322501,	0.077391361,	0.125394398,	0.054844744,	0.202257326)

ks.test(Malates307UWild, Malates307ULab, alternative = "two.sided")
wilcox.test(Malates307UWild, Malates307ULab)


Malates307WWild <- c(0.055515752, 0.632307033, 0.031565495, 0.019285251, 0.030287852, 0.415718807, 0.622797335, 0.031439526, 0.054694469, 0.057132876, 0.409414035, 0.626892572, 0.630405026, 0.632195069, 0.643940884, 0.031146816, 0.039385137, 0.414530055, 0.023882629, 0.413253478, 0.408673239)

Malates307WLab <- c(0.008685914, 0.025398776, 0.005684322, 0.02018242, 0.007460038, 0.009839017, 0.012284117, 0.083390702, 0.014637888, 0.030976206, 0.120827739, 0.060441613, 0.168027062, 0.006032581, 0.163690533, 0.032222558, 0.012776098, 0.02698675, 0.009315799, 0.016589876, 0.008142539, 0.076273366, 0.021305067, 0.024379384, 0.113655396, 0.053495283, 0.173299675, 0.012875471, 0.169043911, 0.020805976, 0.006253476, 0.028541819, 0.016610686, 0.035964186, 0.107915659, 0.014511711, 0.055301292, 0.145303422, 0.085030998, 0.147046853, 0.020444322, 0.142740195, 0.01584782, 0.011127909, 0.005637383, 0.016616325, 0.087957202, 0.010411098, 0.035308077, 0.125334709, 0.065029275, 0.16373194, 0.003179059, 0.159542427, 0.023457074, 0.011174292, 0.030377999, 0.102387655, 0.008482951, 0.049748222, 0.139808238, 0.079463906, 0.151546854, 0.015015883, 0.147226999, 0.013245064, 0.010283493, 0.080593597, 0.01783534, 0.028325309, 0.118218616, 0.057677567, 0.170329648, 0.009872595, 0.166274712, 0.019724404, 0.091663371, 0.005079286, 0.038986197, 0.12910964, 0.068654386, 0.160692678, 0.00441444, 0.156418157, 0.072453784, 0.023837689, 0.021117765, 0.109843501, 0.049572312, 0.176408473, 0.016102586, 0.172136015, 0.095895744, 0.052915841, 0.038527916, 0.026893534, 0.236059666, 0.087798845, 0.231764908, 0.043523264, 0.133289622, 0.072920761, 0.157315979, 0.009255414, 0.152993487, 0.090335089, 0.029993981, 0.192685193, 0.035209254, 0.188402365, 0.063471356, 0.266873657, 0.125192514, 0.262557763, 0.217260566, 0.064772448, 0.212950227, 0.163646185, 0.004991059, 0.159406513)
ks.test(Malates307WWild, Malates307WLab)
wilcox.test(Malates307WWild, Malates307WLab)


Malates270WWild <- c(0.526327927, 0.542581168, 0.086099854, 0.163741843, 0.545132442, 0.553658382, 0.091629261, 0.266740891, 0.018818383, 0.525398339, 0.623788371, 0.021859163, 0.031332345, 0.550093438, 0.294198507, 0.541304565, 0.63900397, 0.005365795, 0.014110164, 0.564708785, 0.311773612, 0.577153764, 0.323915496, 0.291604045, 0.009592332, 0.568669139, 0.314784685,0.641132357, 0.649063615, 0.150265017, 0.37807203, 0.185108683, 0.543803105, 0.552114966, 0.109997308, 0.25879184)
Malates270WLab <- c(0.156521702, 0.295885137, 0.210456794, 0.20754505, 0.261955766, 0.171130775, 0.204498158, 0.306664396, 0.271805154, 0.23956664, 0.146164876, 0.057667508, 0.060195586, 0.111436068, 0.033828484, 0.056989109, 0.437983775, 0.121022657, 0.087787573, 0.098489872, 0.142971363, 0.035675843, 0.131821921, 0.097448805, 0.556385054, 0.026168639, 0.060022376, 0.061268211, 0.063488976, 0.046919256, 0.02093133, 0.482576404, 0.073163423, 0.041967429, 0.109344495, 0.059244951, 0.06888997, 0.490397496, 0.11858302, 0.087140209, 0.096903536, 0.062313682, 0.525313959, 0.009777743, 0.025515803, 0.535199319, 0.505767763, 0.034316108, 0.469961728, 0.071954865, 0.038492903, 0.03694498, 0.439331887, 0.106481829, 0.073118943)

ks.test(Malates270WWild, Malates270WLab)
wilcox.test(Malates270WLab, Malates270WWild)


Malates270UWild <- c(0.76630486, 0.731697931, 0.512614894, 0.661144985, 0.817974547, 0.839378469, 0.563368056, 0.56078063, 0.434852259, 0.750397699, 0.856996141, 0.333720752, 0.37400244, 0.732635629, 0.728078739, 0.715678461, 0.836667326, 0.413790117, 0.465319505, 0.688258047, 0.641577938, 0.629361907, 0.802452899, 0.825327041, 0.475598176, 0.518452238, 0.821047874, 0.788718076, 0.534220649, 0.284897811, 0.796610578, 0.733753189, 0.873767603, 0.900140834, 0.605250277, 0.675435885)
Malates270ULab <- c(0.324522575, 0.482821429, 0.291348582, 0.216298678, 0.433905586, 0.386768655, 0.351142214, 0.387696427, 0.533078416, 0.434163778, 0.53050892, 0.329132402, 0.296180275, 0.465685894, 0.452895784, 0.454780553, 0.491380282, 0.564694755, 0.471396203, 0.393848013, 0.411568729, 0.307572341, 0.336703766, 0.282132107, 0.464750839, 0.180477728, 0.284403022, 0.248779561, 0.316269804, 0.375386473, 0.353188919, 0.398315928, 0.439840014, 0.3554125, 0.372441852, 0.39038056, 0.369759719, 0.42982748, 0.481252836, 0.404324643, 0.479410838, 0.385972973, 0.232465504, 0.292734045, 0.313359082, 0.172560147, 0.154286383, 0.332392678, 0.35019787, 0.233789846, 0.256792702, 0.2012755, 0.372566078, 0.200868369, 0.156479815)
ks.test(Malates270UWild, Malates270ULab)
wilcox.test(Malates270UWild, Malates270ULab)


Workers307WWild <- c(0.112266307, 0.142590166, 0.21604023, 0.090348913, 0.281901703, 0.326720287, 0.095649144, 0.208136751, 0.054048416, 0.286981444, 0.326329023, 0.274542078, 0.316273015, 0.112037654, 0.184444402, 0.115810404, 0.179128637, 0.143983782, 0.09484676, 0.214611651, 0.23721783)
Workers307WLab <- c(0.109104083, 0.096003683, 0.094464282, 0.066483783, 0.104056923, 0.161018923, 0.171690594, 0.022436715, 0.084023984, 0.093859757, 0.09730098, 0.140354693, 0.184823863, 0.190456148, 0.116188587, 0.196695966, 0.059401681, 0.231307763, 0.119912339, 0.187628031, 0.189536433, 0.188584334, 0.070660956, 0.02446539, 0.154415472, 0.033865524, 0.238685613, 0.154089624, 0.095805882, 0.018390989, 0.049077584, 0.049904285, 0.230529531, 0.136858658, 0.015902744, 0.240623891, 0.135545492, 0.077615595,0.014236957, 0.029988164, 0.031561666, 0.213745682, 0.1305297, 0.159406315, 0.125871155, 0.06426246, 0.142195285, 0.112621388, 0.112639887, 0.095559869, 0.243124757, 0.131750868, 0.084164662, 0.024555078, 0.03218963, 0.030339165, 0.209480214, 0.272497138, 0.166557059, 0.24030006, 0.241023788, 0.242467866, 0.114339497, 0.160857082, 0.143669057, 0.115242678, 0.112707868, 0.194277345, 0.015153912, 0.18919185, 0.190951444, 0.034826092, 0.036105138, 0.217744718, 0.081312588, 0.076336752, 0.080968805, 0.139271007)
ks.tet(Workers307WLab, Workers307WWild)
wilcox.test(Workers307WLab, Workers307WWild)
Workers307UWild <- c(0.485970046, 0.388193073, 0.377299974, 0.419584461, 0.286692786, 0.303280424, 0.351400575, 0.393763452, 0.429762649, 0.424176356, 0.451787049, 0.379735856, 0.405006557, 0.313691745, 0.312057814, 0.353465926, 0.270633623, 0.26635648, 0.349389034, 0.306843025, 0.16795583)
Workers307ULab <- c(0.27773311, 0.246363941, 0.176211112, 0.374607003, 0.211674804, 0.174997789, 0.424136027, 0.175182376, 0.262581941, 0.502334298, 0.305776963, 0.350758068, 0.155884993, 0.298594778, 0.287901686, 0.323414529, 0.288467581, 0.360164521, 0.19896106, 0.379435124, 0.544055321, 0.321038311, 0.307119889, 0.203942179, 0.324122155, 0.257786149, 0.2712786, 0.336515694, 0.111160657, 0.315179544, 0.504685114, 0.274029284, 0.300885109, 0.390776514, 0.170591101, 0.136006877, 0.426466157, 0.16399974, 0.253292228, 0.500441828, 0.295804645, 0.323204753, 0.400875118, 0.388385626, 0.260486365, 0.364356918, 0.439738346, 0.510950473, 0.36402698, 0.218489371, 0.219669546, 0.453866033, 0.190882588, 0.299703288, 0.509819028, 0.310633141, 0.374361072, 0.435366305, 0.195623265, 0.292119997, 0.538688993, 0.332402036, 0.305725605, 0.372471537, 0.471765088, 0.544696292, 0.415930299, 0.299714065, 0.423193111, 0.449742749, 0.38267575, 0.493892647, 0.359862389, 0.422230995, 0.278070207, 0.490633989, 0.252002974, 0.341307596)
ks.test(Workers307UWild, Workers307ULab)
t.test(Workers307UWild, Workers307ULab)

Workers270WWild <- c(0.069928698, 0.413524927, 0.100013405, 0.211387094, 0.328655728, 0.193826176, 0.197163325, 0.197194809, 0.491757159, 0.079800773, 0.395360124, 0.425602845, 0.102978055, 0.205475411, 0.340227703, 0.213430121, 0.213378435, 0.179181941, 0.502994097, 0.079430754, 0.407027955, 0.396417966, 	0.258220922, 0.151146449, 0.261798813, 0.253821955, 0.29276643, 0.179216855, 0.393471377, 0.073381097, 0.232138665, 0.348620522, 0.230705028, 0.197733452, 0.223249405, 0.498164727, 0.093719237, 0.37872449, 0.152035084, 0.086803538, 	0.063654338, 0.05364936, 0.343370341, 0.1795834, 0.237453255, 0.161415191, 0.16127523, 0.188095513, 0.213911615, 0.311532312, 0.22794077, 0.08609219, 0.091786886, 0.349494486, 0.201968447, 0.237287888, 0.09689261, 0.337094224, 0.152967831, 0.232969234, 0.471575742, 0.24777716, 0.374534671, 0.370452194, 0.155078988, 0.272737011)
Workers270WLab <- c(0.111470807, 0.136396094, 0.091482419, 0.076320702, 0.089565355, 0.048444957, 0.053823764, 0.058791391, 0.087128527, 0.097884941, 0.064915856, 0.092627254, 0.089918308, 0.170689233, 0.105747673, 0.141203034, 0.153529072, 0.163277152, 0.069878007, 0.182347613, 0.085665433, 0.105238983, 0.180557346, 0.094988883, 0.162845885, 0.164921371, 0.166975005, 0.103028312, 0.204816669, 0.134868764, 0.114281242, 0.035518629, 0.085428123, 0.086082269, 0.118847738, 0.032727071, 0.14227685, 0.077475398, 0.112342218, 0.053239001, 0.051937587, 0.033947536, 0.112842088, 0.039497318, 0.106327122, 0.102412449, 0.094260873, 0.120079975, 0.045326678, 0.145624947, 0.087947477, 0.01954538, 0.056310399, 0.084529033, 0.067108572, 0.088269003, 0.13274919, 0.064431373, 0.104980993, 0.11087899, 0.049854378, 0.098125615, 0.06071593, 0.091696423, 0.071529594, 0.098926807)
ks.test(Workers270WWild, Workers270WLab)
wilcox.test(Workers270WWild, Workers270WLab)


Workers270UWild <- c(0.574, 0.576, 0.555, 0.541, 0.610, 0.534, 0.599, 0.541, 0.749, 0.455, 0.652, 0.512, 0.425, 0.465, 0.411, 0.505, 0.440, 0.478, 0.642, 0.437, 0.480, 0.423, 0.300, 0.358, 0.528, 0.417, 0.445, 0.523, 0.527, 0.470, 0.341, 0.388, 0.554, 0.394, 0.406, 0.607, 0.456, 0.516, 0.304, 0.483, 0.401, 0.370, 0.533, 0.478, 0.395, 0.546, 0.417, 0.397, 0.517, 0.478, 0.369, 0.495, 0.562, 0.722, 0.489, 0.568, 0.733, 0.516, 0.543, 0.538, 0.572, 0.505, 0.452, 0.615, 0.481, 0.538)
Workers270ULab <- c(0.222, 0.291, 0.213, 0.255, 0.253, 0.293, 0.275, 0.145, 0.275, 0.213, 0.212, 0.294, 0.186, 0.107, 0.237, 0.174, 0.228, 0.154, 0.281, 0.171, 0.121, 0.298, 0.293, 0.169, 0.340, 0.238, 0.262, 0.213, 0.270, 0.216, 0.198, 0.219, 0.280, 0.137, 0.189, 0.244, 0.272, 0.260, 0.277, 0.180, 0.259, 0.189, 0.255, 0.189, 0.121, 0.310, 0.154, 0.191, 0.190, 0.248, 0.243, 0.302, 0.229, 0.316, 0.244, 0.195, 0.278, 0.258, 0.086, 0.260, 0.230, 0.192, 0.217, 0.156, 0.331, 0.284)
ks.test(Workers270UWild, Workers270ULab)
wilcox.test(Workers270UWild, Workers270ULab)
```
