## Supplemental Figures and Tables for "*Trachymyrmex septentrionalis* ant microbiome assembly is unique to individual colonies and castes"

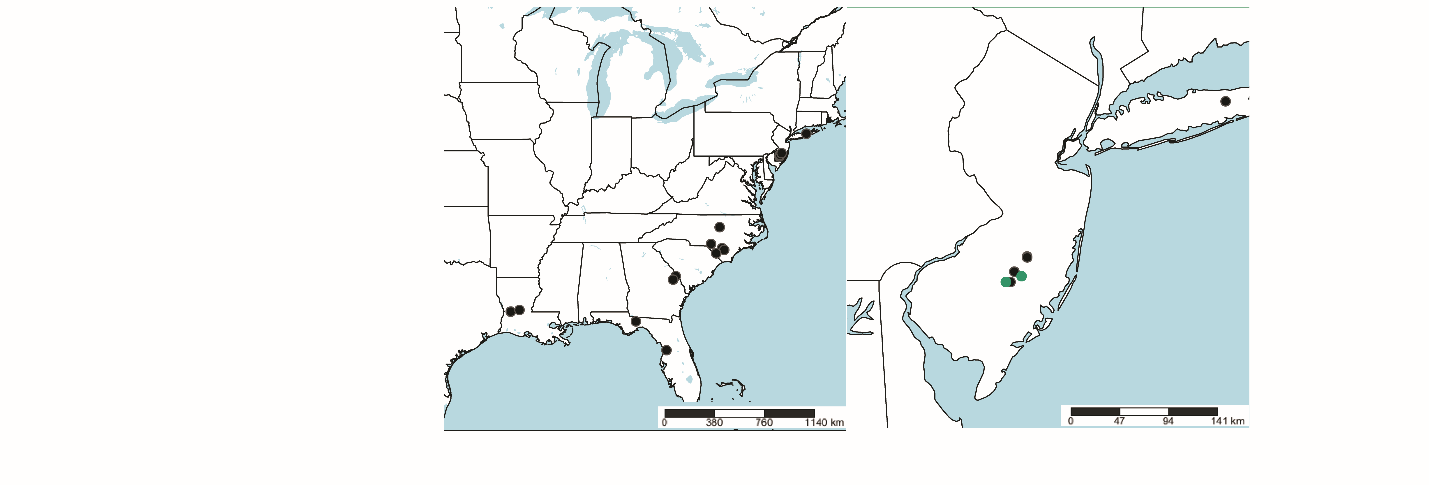

Supplementary Figure S1) Maps detailing colony locations. The left map shows the location of the colonies in the multi-state dataset and two-colony dataset using black circles. On the right, New Jersey is magnified to show colonies from the multi-state dataset (black circles) and the two-colony dataset (green circles). Maps were created using www.simplemappr.net/ and edited in Adobe Illustrator.

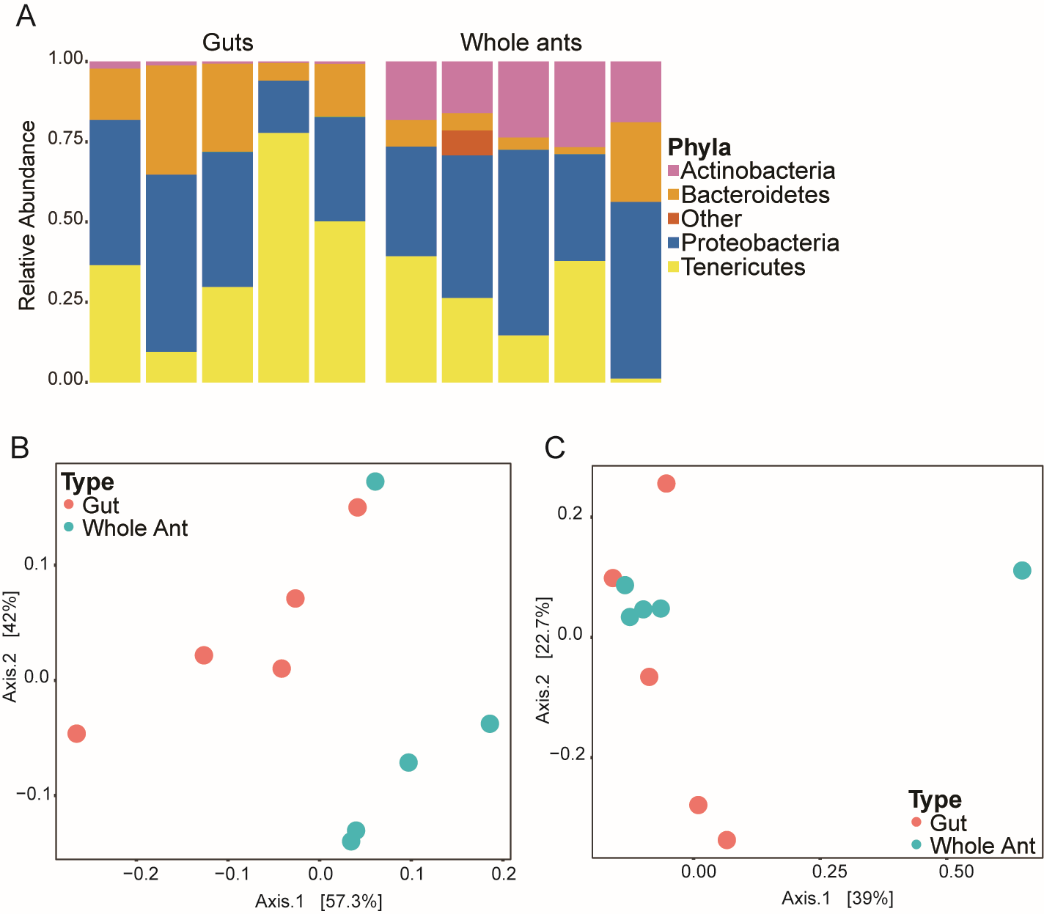

Supplementary Figure S2) Comparison of whole ant and gut microbiomes. A) The phyla present in whole ant and ant gut microbiomes at > 15% abundance. Single bars represent individual ants. The phyla are differentiated by color with “Other” representing phyla present at < 15% relative abundance. B) and C): PCoA ordinations of Weighted (B) and Unweighted (C) Unifrac distances between whole ant and ant gut microbiomes. Sample types are indicated using color. n=10. WUF PERMANOVA: R^2^=0.342, p= 0.01; UUF PERMANOVA: R^2^=0.110, p= 0.451; Bray-Curtis R^2^=0.352, p= 0.053.

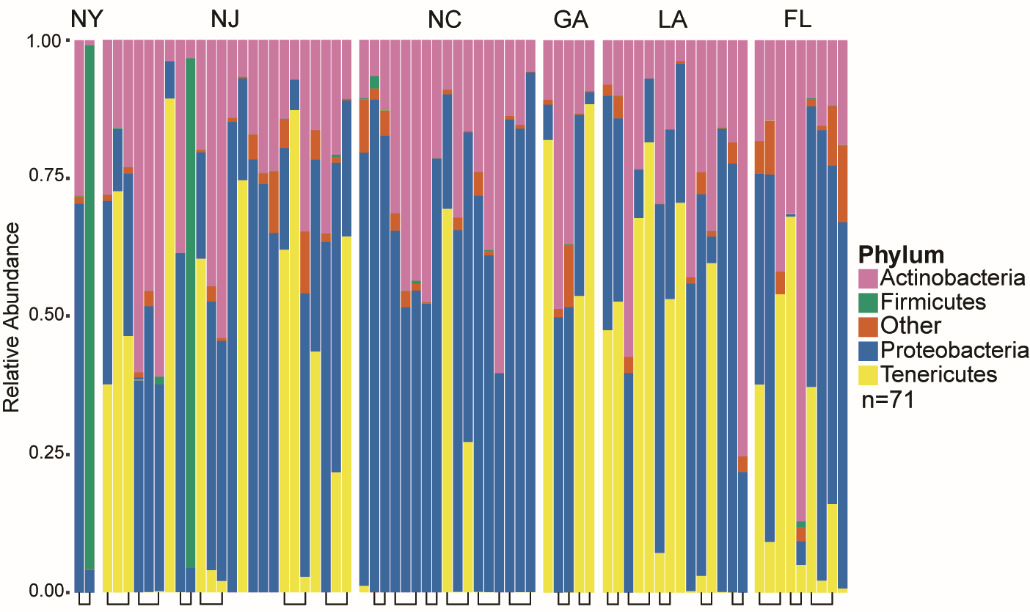

Supplementary Figure S3) Bar plot of the phyla found in samples from the multi-state dataset. Single bars represent a single ant microbiome and are grouped by state of collection. X-axis backets group worker ants from the same colony. Phyla are differentiated by color, “Other” represents Phyla present at >15%. n=71

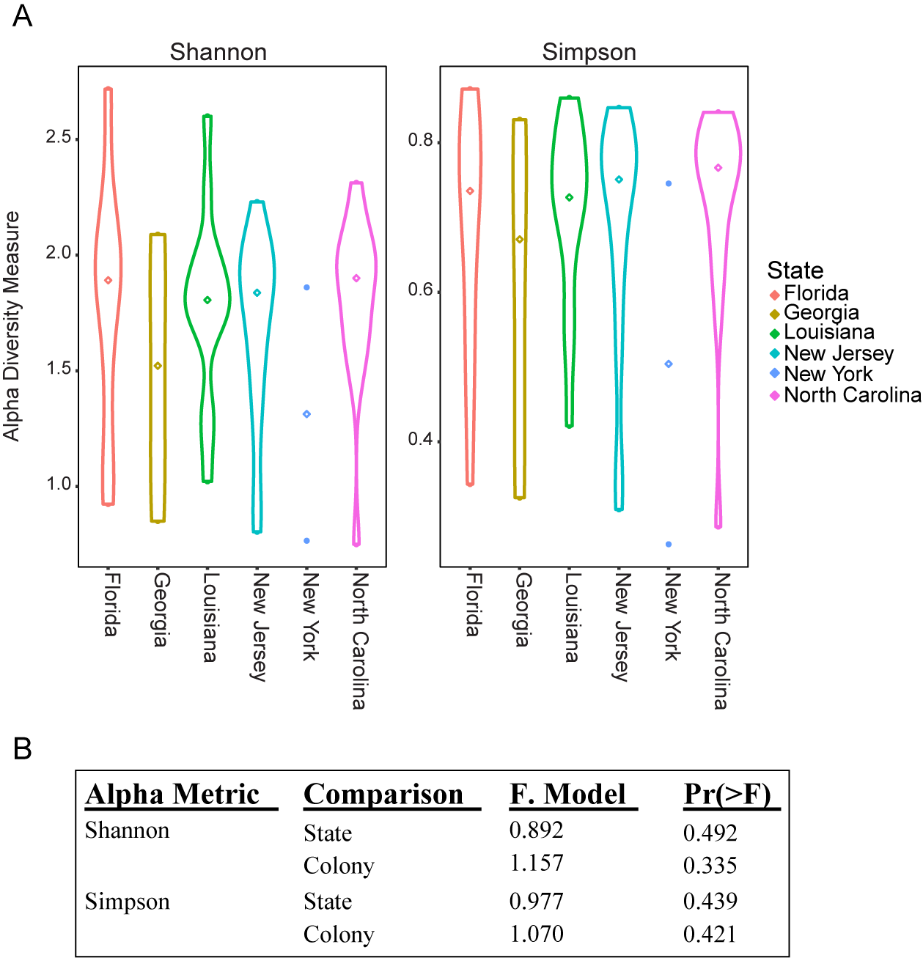

Supplementary Figure S4) A) Shannon and Simpson alpha-diversities for samples in the multi-state dataset. Median α-diversities are marked with diamonds. B) ANOVA of alpha diversity scores for the multistate dataset, compared between states and colonies. n=71

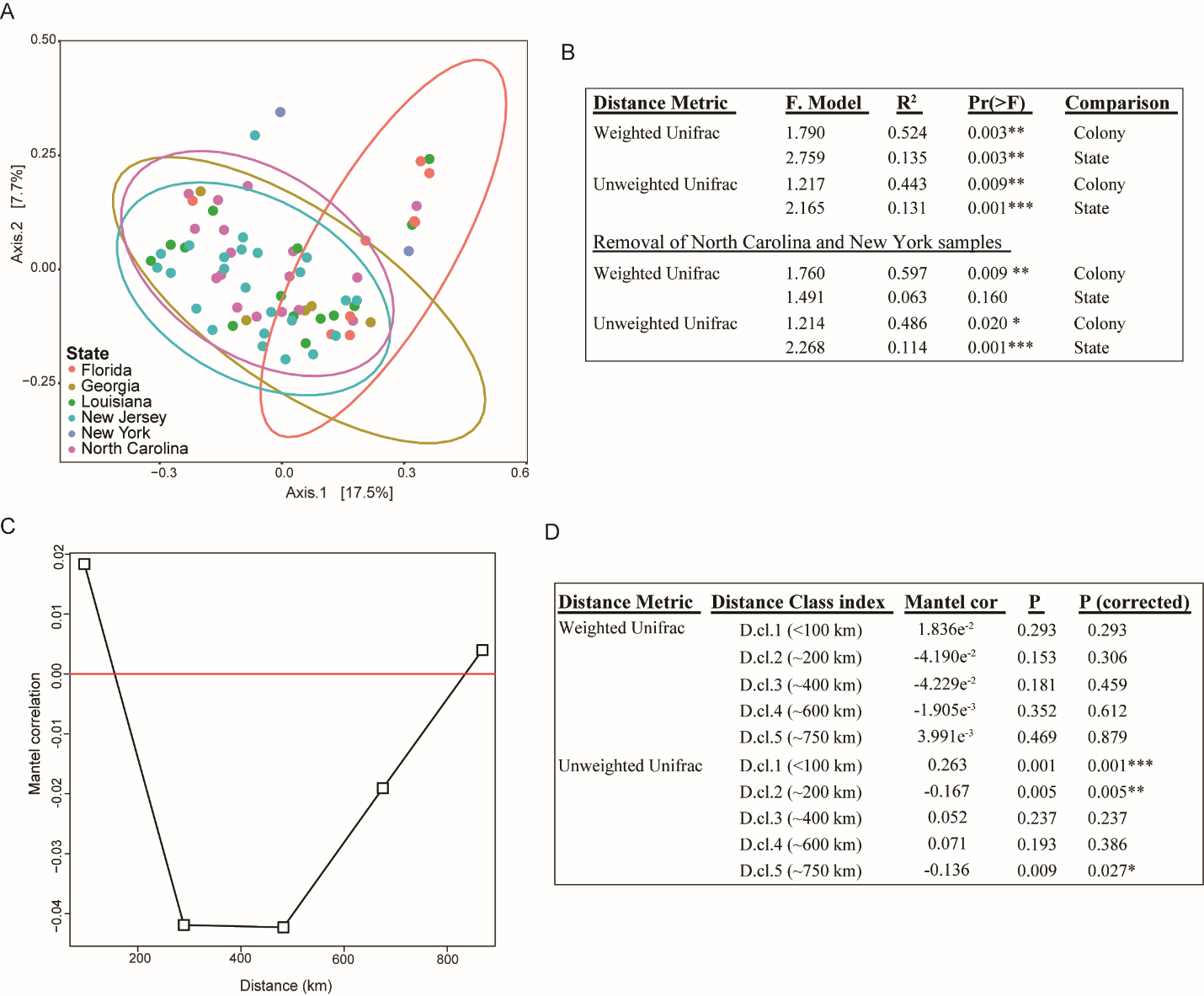

Supplemental Figure S5) A) PCoA of unweighted Unifrac distances between ant microbiomes in the multi-state dataset. Samples are colored by state and grouped using a multivariate t-distribution. B) PERMANOVA analyses testing the variation of ant microbiomes in the multi-state dataset explained by state of collection (New York, New Jersey, North Carolina, Georgia, Florida, and Louisiana) or colony of origin. The bottom statistics exclude North Carolina and New Jersey, which lacked Tenericute symbionts. C) Mantel correlation plot of weighted Unifrac distances and ant colony geographic location (GPS points) of the multi-state dataset. The x-axis is distance class index in kilometers, and y-axis the Mantel correlation R statistic. Filled squares indicate significant p values (p < 0.05), and unfilled squares indicate nonsignificant p values (p > 0.05). Points above and below the red line indicate positive and negative correlations, respectively. D) Partial Mantel statistic scores for the comparison of microbiome similarity and geographic distance in the multistate dataset. For B & C) Significance codes: ‘***’≥ 0.001, ‘**’≥ 0.01, ‘*’ ≥0.05.

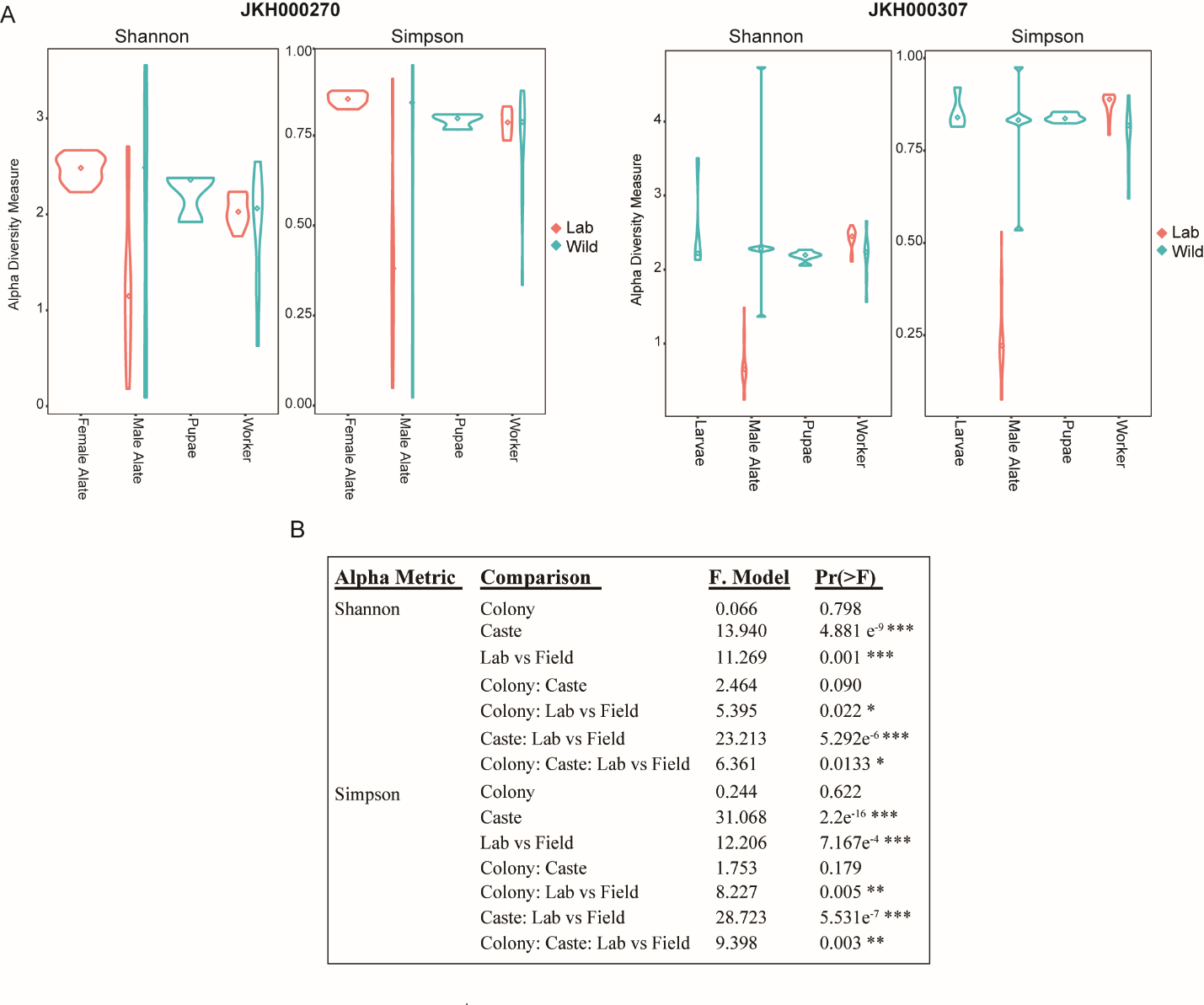

Supplementary Figure S6) A) Shannon and Simpson alpha-diversities for colonies JKH000270 and JKH000307. The x-axes list each caste, and median α-diversities are marked with diamonds. B) ANOVA of alpha diversity scores for the two-colony dataset, compared between colonies JKH000270 & JKH000307, caste, and whether ants were lab-maintained or sampled in the field. Significance codes: ‘***’≥ 0.001, ‘**’≥ 0.01, ‘*’ ≥0.05

n=110

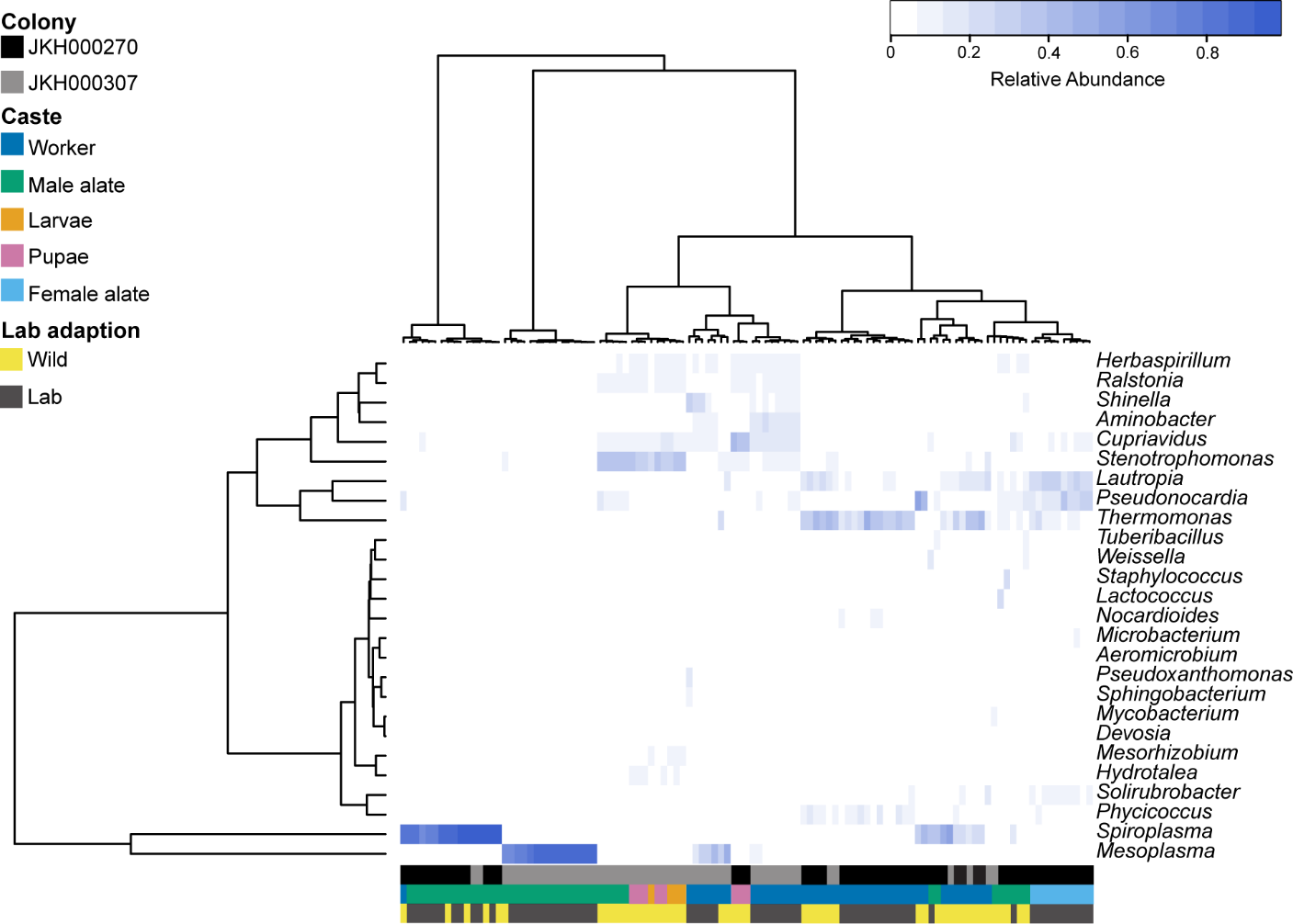

Supplementary Figure S7) Heatmap of bacterial genera in the two colonies dataset with an abundance of ≥ 5% in at least one sample. On the left, a Euclidian distance dendrogram represents the relationship between the relative abundance of reads in each genus, labeled on the right. The top dendrogram clusters the ant microbiomes using the ward.D distance. On the bottom, colors in the three different rows differentiate samples by their colony, caste, and whether they are lab-adapted. n=110

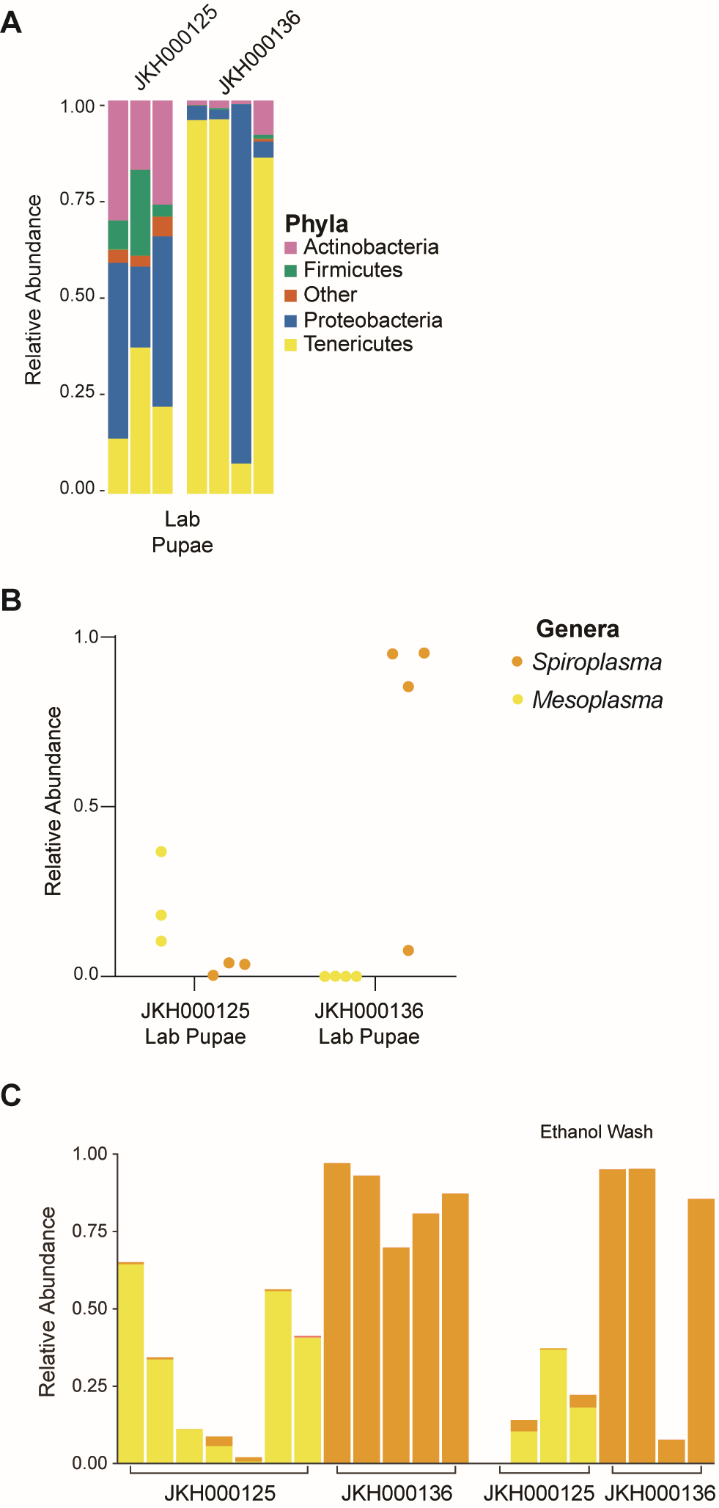

Supplementary Figure S8) A) Bar plot of the top phyla in pupae from colony JKH000125 (left) and JKH000136 (right). Phyla are differentiated by color and “Other” represents phyla present at < 15% relative abundance. B) *Spiroplasma* and *Mesoplasma* relative abundance in pupae from colony JKH000136 and JKH000125. C) *Mesoplasma* and *Spiroplasma* relative abundances in pupae from colonies JKH000125 and JKH000136. Single bars represent abundances in single pupae and x-axis brackets group pupae from each colony. Samples shown to the right of the figure were ethanol washed before DNA extraction, unlike those on the left.

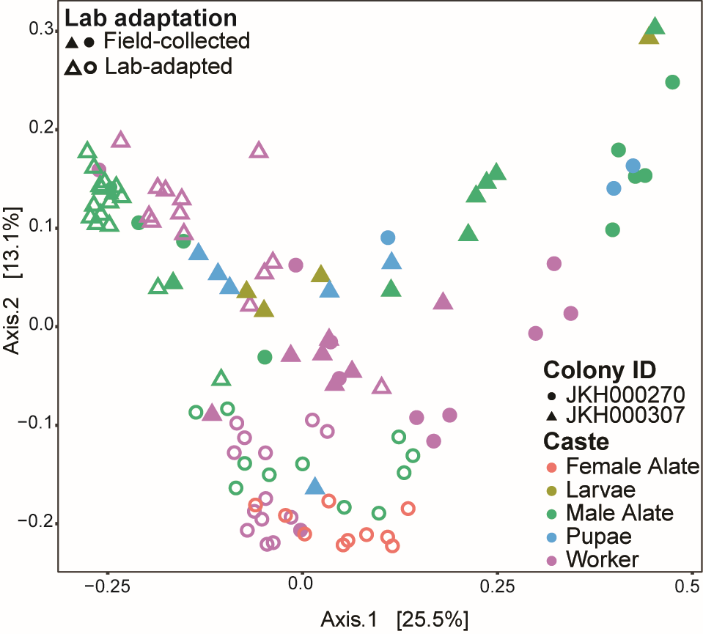

Supplementary Figure S9) PCoA of Unweighted Unifrac distances between ant microbiomes in the two-colony dataset. Colors indicate ant caste, shape indicates colony ID, and solid and open shapes indicate field-collected and lab-maintained samples, respectively. n= 110

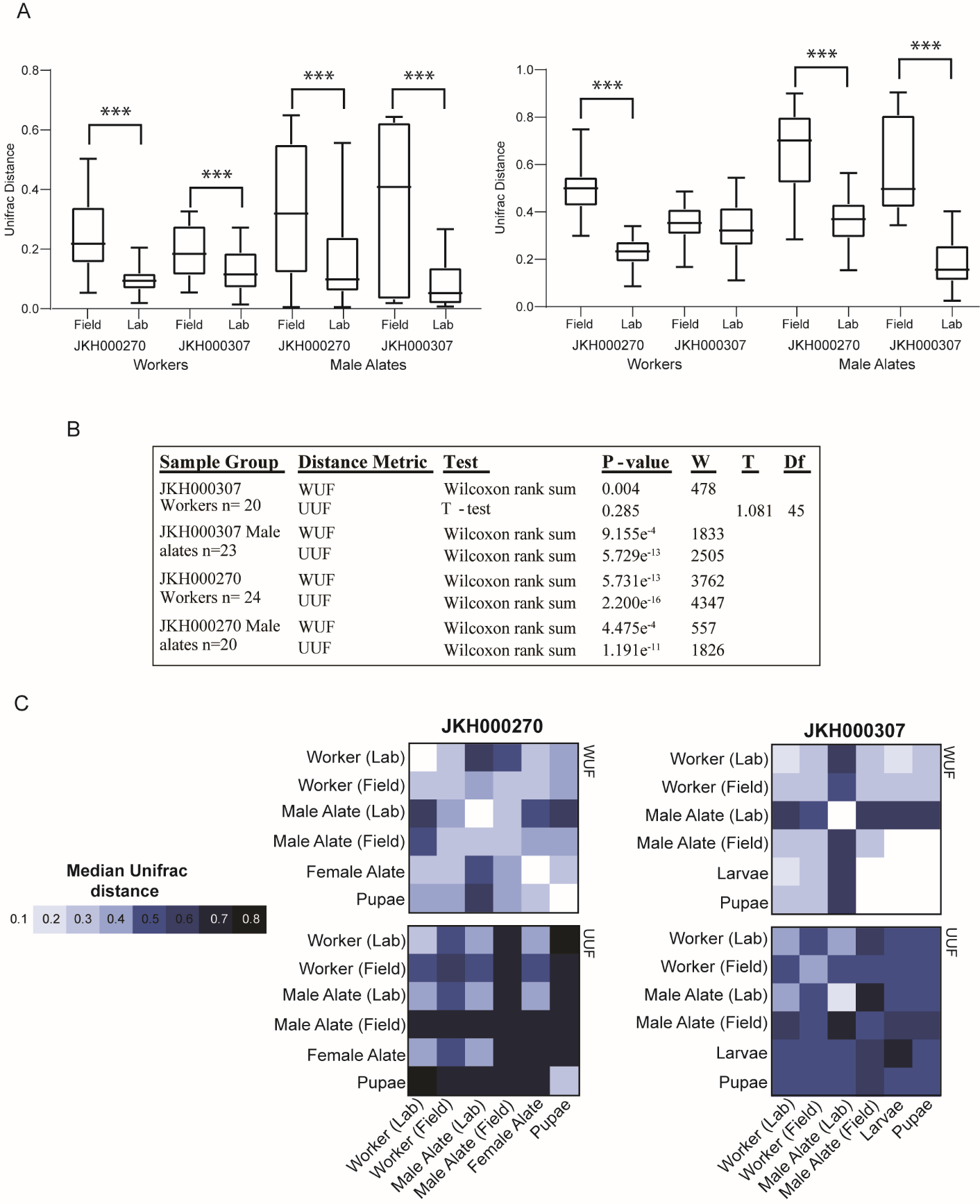

Supplementary Figure S10) A) Box and whisker plots comparing distributions of weighted (right) and unweighted (left) Unifrac distances for lab-maintained and field-collected samples from each caste and colony. Boxes represent the upper and lower quartiles, whiskers represent the maximum and minimum values, and the middle bar represents the median Brackets between two samples indicate a difference of statistical significance. Significance codes: ‘***’≥ 0.001. B) Analyses testing if the variation in weighted or unweighted Unifrac distances differ between lab-maintained and field-collected microbiomes, comparing only castes and colonies for which both lab-maintained and field-collected samples were available. C) Median weighted (top) and unweighted (bottom) Unifrac distances within and between sample samples from the same caste and lab-adaptation categories for each colony. Colors indicate median Unifrac distances between samples belonging to each category.
